# The differentiation and integration of the hippocampal dorsoventral axis are controlled by two nuclear receptor genes

**DOI:** 10.1101/2023.02.17.528915

**Authors:** Xiong Yang, Rong Wan, Zhiwen Liu, Su Feng, Jiaxin Yang, Naihe Jing, Ke Tang

## Abstract

The hippocampus executes crucial functions from declarative memory to adaptive behaviors associated with cognition and emotion. However, the mechanisms of how morphogenesis and functions along the hippocampal dorsoventral axis are differentiated and integrated are still largely unclear. Here, we show that *Nr2f1* and *Nr2f2* genes are distinctively expressed in the dorsal and ventral hippocampus, respectively. The loss of *Nr2f2* results in ectopic CA1/CA3 domains in the ventral hippocampus. The deficiency of *Nr2f1* leads to the failed specification of dorsal CA1, among which there are place cells. The deletion of both *Nr2f* genes causes almost agenesis of the hippocampus with abnormalities of trisynaptic circuit and adult neurogenesis. Moreover, *Nr2f1/2* may cooperate to guarantee appropriate morphogenesis and function of the hippocampus by regulating the *Lhx5-Lhx2* axis. Our findings revealed a novel mechanism that *Nr2f1* and *Nr2f2* converge to govern the differentiation and integration of distinct characteristics of the hippocampus in mice.

## Introduction

Memory, including declarative and nondeclarative memory, unifies our mental world to ensure the quality of life for people of all ages, from newborns to elderly individuals (Eichenbaum & Cohen, 2014; Kandel, Dudai, & Mayford, 2014). The pioneering studies of Milner and her colleagues revealed that the hippocampus is required for declarative memory but not nondeclarative memory (Penfield & Milner, 1958; Scoville & Milner, 1957). The discovery of activity-dependent long-term potentiation and place cells provides the neurophysiological basis of hippocampal function (Bliss & Lomo, 1973; O’Keefe & Dostrovsky, 1971). The rodent hippocampus can be divided into the dorsal and ventral domains, corresponding to the posterior and anterior hippocampus in humans, respectively. In recent decades, numerous studies have supported the Moser theory that the hippocampus is a heterogeneous structure with distinct characteristics of gene expression, connectivity, and functions along its dorsoventral axis (Bast, 2007; Fanselow & Dong, 2010; Moser & Moser, 1998; Strange, Witter, Lein, & Moser, 2014). The dorsal hippocampus, which connects and shares similar gene expression with the neocortex (Fanselow & Dong, 2010), serves the “cold” cognitive function associated with declarative memory and spatial navigation. The ventral hippocampus, which connects and generates similar gene expression with the amygdala and hypothalamus (Cenquizca & Swanson, 2007; Kishi et al., 2000; Pitkanen, Pikkarainen, Nurminen, & Ylinen, 2000), corresponds to the “hot” affective states related to emotion and anxiety (Fanselow & Dong, 2010; Tyng, Amin, Saad, & Malik, 2017). Nonetheless, to date, the molecular and cellular mechanisms by which the morphogenesis, connectivity, and functions along the dorsoventral axis of the hippocampus are differentiated and integrated are largely unknown.

The hippocampus, a medial temporal lobe structure in the adult rodent forebrain, originates from the medial pallium (MP) in the medial line of the early dorsal telencephalon. The cortical hem (CH), which is located ventrally to the MP, functions as an organizer for hippocampal development (Hebert & Fishell, 2008; Schuurmans & Guillemot, 2002). It has been demonstrated that both extrinsic signals, such as WNTs and BMPs, and intrinsic factors, including EMX1, EMX2, LEF1, LHX2, and LHX5, are involved in the regulation of early morphogenesis of the hippocampus. As the earliest *Wnt* gene to be exclusively expressed in the cortical hem, *Wnt3a* is required for the genesis of the hippocampus (S. M. Lee, Tole, Grove, & McMahon, 2000); in addition, *Lef1* is downstream of Wnt signaling, and the hippocampus is completely absent in *Lef1^neo/neo^* null mutant mice (Galceran, Miyashita-Lin, Devaney, Rubenstein, & Grosschedl, 2000). Wnt signaling is essential for early development of the hippocampus. *Emx1* and *Emx2* are mouse homologs of *Drosophila empty spiracles* (Simeone et al., 1992). Interestingly, the dorsal hippocampus is smaller in an *Emx1* null mutant (Yoshida et al., 1997), while *Emx2* is required for the growth of the hippocampus but not for the specification of hippocampal lineages (Tole, Goudreau, Assimacopoulos, & Grove, 2000). Moreover, *Lhx5*, which encodes a LIM homeobox transcription factor and is specifically expressed in the hippocampal primordium, is necessary for the formation of the hippocampus (Zhao et al., 1999). *Lhx2*, encoding another LIM homeobox transcription factor, is required for the development of both the hippocampus and neocortex (Mangale et al., 2008; Monuki, Porter, & Walsh, 2001; Porter et al., 1997). Intriguingly, deficiency of either *Lhx5* or *Lhx2* results in agenesis of the hippocampus, and more particularly, these genes inhibit each other (Hebert & Fishell, 2008; Mangale et al., 2008; Roy, Gonzalez-Gomez, Pierani, Meyer, & Tole, 2014; Zhao et al., 1999), indicating that the *Lhx5* and *Lhx2* genes may generate an essential regulatory axis to ensure the appropriate hippocampal development. Nevertheless, whether there are other intrinsic genes that participate in the regulation of morphogenesis and function of the hippocampus has not been fully elucidated.

*Nr2f* genes, including *Nr2f1* and *Nr2f2,* encode two transcription factor proteins belonging to the nuclear receptor superfamily (Yang, Feng, & Tang, 2017). Mutations of *Nr2f1* are highly related to neurodevelopmental disorders (NDD), such as intellectual disability (ID) and autism spectrum disorders (ASD) (Bertacchi et al., 2020; Bosch et al., 2014; Contesse, Ayrault, Mantegazza, Studer, & Deschaux, 2019), and mutations of the *Nr2f2* gene are associated with congenital heart defects (CHD) (Al Turki et al., 2014). By using animal models, our studies and others have demonstrated that *Nr2f* genes participate in the regulation of the development of the central nervous system (Zhang et al., 2020). The *Nr2f1* plays an essential role in the differentiation of cortical excitatory projection neurons and inhibitory interneurons, the development of the dorsal hippocampus, and cortical arealization (Armentano et al., 2007; Bertacchi et al., 2020; Del Pino et al., 2020; J. Feng et al., 2021; Flore et al., 2017; Lodato et al., 2011; C. Zhou et al., 1999; C. Zhou, Tsai, & Tsai, 2001). *Nr2f2* plays a vital role in the development of the amygdala, hypothalamus, and cerebellum (S. Feng et al., 2017; Kim, Takamoto, Yan, Tsai, & Tsai, 2009; Tang, Rubenstein, Tsai, & Tsai, 2012). Nevertheless, whether and how *Nr2f1* and/or *Nr2f2* genes regulate the differentiation and integration of hippocampal morphogenesis, connectivity, and function is still largely unclear.

Here, our data show that *Nr2f1* and *Nr2f2* genes are differentially expressed along the dorsoventral axis of the postnatal hippocampus. The loss of *Nr2f2* results in ectopic CA1 and CA3 domains in the ventral hippocampus. In addition, the deficiency of *Nr2f1* leads to not only dysplasia of the dorsal hippocampus but also failed specification and differentiation of the dorsal CA1 pyramidal neuron lineage. Furthermore, the deletion of both genes in the RX^Cre/+^; *Nr2f1^F/F^*; *Nr2f2^F/F^* double mutant mouse causes almost agenesis of the hippocampus, accompanied by compromised specification of the CA1, CA3, and dentate gyrus (DG) domains. The components of the trisynaptic circuit are abnormal in the corresponding single-gene or double-gene mutant model. Moreover, *Nr2f* genes may cooperate to guarantee the appropriate morphogenesis and function of the hippocampus by regulating the *Lhx5-Lhx2* axis.

## Results

### Differential expression profiles of *Nr2f1* and *Nr2f2* genes along the dorsoventral axis in the developing and postnatal hippocampus

To investigate the functions of the *Nr2f1* and *Nr2f2* genes in the hippocampus, immunofluorescence staining was first performed to examine their expression in wild-type mice at postnatal month 1 (1M). In both the coronal and sagittal sections, NR2F1 exhibited a septal/dorsal high-temporal/ventral low expression pattern along the hippocampus (Figure 1Aa, d, c, f, g, j, m, i, l, and o), and its expression is highest in the dorsal CA1 region (Figure 1Aa, d, c, f, j, and l), where place cells are mainly located (O’Keefe & Conway, 1978; O’Keefe & Dostrovsky, 1971), and the dorsal DG, where there are adult neural stem cells (NSCs) (Gould & Cameron, 1996). However, the expression of NR2F2 was high in the temporal/ventral hippocampus but was barely detected in the septal/dorsal part of the hippocampus (Figure 1Ab, c, e, f, h, i, k, l, n, and o). The dorsal-high NR2F1 and ventral-high NR2F2 expression profiles were further verified in the postnatal hippocampi at 1M by western blotting assays (Figure 1Ap, q). At embryonic day 10.5 (E10.5), NR2F1 was detected in the dorsal pallium (DP) laterally and NR2F2 was expressed in the MP and CH medially (Figure 1—figure supplement 1Aa-b). At E11.5 and E12.5, the expression of NR2F2 remained in the CH (Figure 1—figure supplement 1Ac-d, Bb-c). Interestingly, NR2F1 and NR2F2 generated complementary expression patterns in the hippocampal primordium with NR2F1 in the dorsal MP and NR2F2 in the ventral CH at E14.5 (Figure 1—figure supplement 1Ba-f). Additionally, septal/dorsal-high NR2F1 and temporal/ventral-high NR2F2 expression patterns were observed at postnatal day 0 (P0) (Figure 1—figure supplement 1Bg-l). The data above revealed that the differential expression patterns of *Nr2f1* and *Nr2f2* genes were generated and maintained along the dorsoventral axis in the early hippocampal primordium, the developing and postnatal hippocampus, indicating that they could play distinct roles in the mediation of the morphogenesis and functions of the hippocampus.

**Figure 1.**
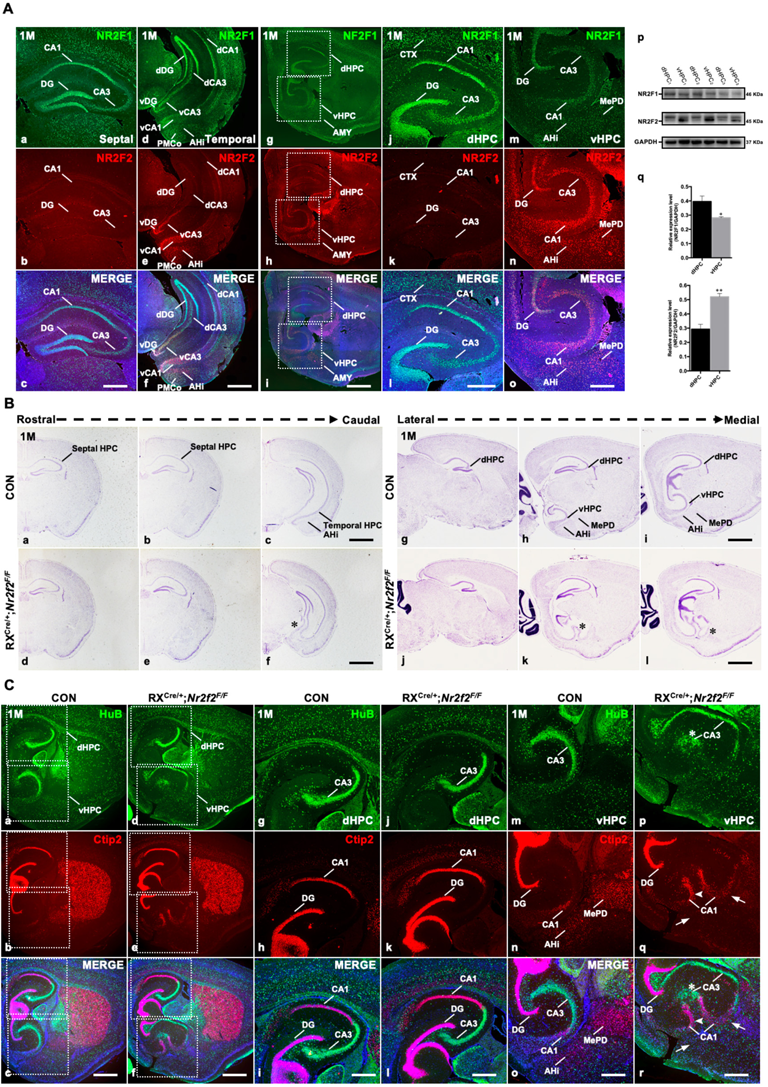
Duplicated CA1 and CA3 domains are generated in the ventral hippocampus of RX^Cre/+^; *Nr2f2^F/F^*mutant mice. A, The expression of NR2F1 (**a, d, g, j, m**) and NR2F2 (**b, e, h, k, n**) in coronal sections (**a-f**) and sagittal sections (**g-o**) of the hippocampus at postnatal month 1 (1M); representative Western blots and quantitative densitometry data for the expression of NR2F1 and NR2F2 in the dorsal and ventral hippocampus at 1M (**p-q**). **B,** In coronal sections along the rostrocaudal axis (**a-f**) and sagittal sections along the lateral-medial axis (**g-l**) of the hippocampus in mutant mice, compared with that in control mice (**a-c**, **g-i**), the ectopic CA-like structure, indicated by the star, was observed in the ventral region in *Nr2f2* gene mutant (RX^Cre/+^; *Nr2f2^F/F^)* mice at 1M (**d-f**, **j-l**). **C,** The expression of HuB and Ctip2 in the corresponding inserted area in **a-f** under a high magnification objective lens at 1M (**g-r**); compared with those of control mice (**a-c**, **g-i**, **m-o**), the duplicated HuB-positive CA3 domain, indicated by the star, and Ctip2-positive domains, indicated by the arrowhead, were specifically observed in the ventral hippocampus (**d-f, p-r**) but not in the dorsal hippocampus (**d-f**, **j-l**) of *Nr2f2* mutant mice at 1M; Ctip2 positive AHi and MePD amygdaloid nuclei were barely observed in the *Nr2f2* mutant mice, indicated by the arrows, instead of the ectopic CA domains at the prospective amygdaloid regions (**e-f**, **q-r**). Data are expressed as the mean ± SEM. *Student’s t test*, \**P*<0.05, \*\**P*<0.01. AHi, amygdalohippocampal area; AMY, amygdala nuclei; CTX, cortex; dCA1, dorsal CA1; dCA3, dorsal CA3; dDG, dorsal dentate gyrus; dHPC, dorsal hippocampus; MePD, posterodorsal part of the medial amygdaloid nucleus; PMCo, posteromedial cortical amygdaloid nucleus; vCA1, ventral CA1; vCA3, ventral CA3; vDG, ventral dentate gyrus; vHPC, ventral hippocampus. Scale bars, **Aa-c**, **Ad-f**, **Aj-o**, **Cg-r**, 100 μm; **Ag-i**, **Ba-l, Ca-f**, 200μm.

Next, to investigate the roles of *Nr2f* genes in the hippocampus, an RX^Cre^ mouse was used to excise the expression of the *Nr2f1* and/or *Nr2f2* genes (Swindell et al., 2006; Tang et al., 2012). The deletion efficiency of RXCre recombinase was verified by immunofluorescence assays. Compared with control mice, either *Nr2f2* or *Nr2f1* could be excised in the postnatal hippocampus of corresponding single-gene mutants at 1M (Figure 1—figure supplement 1Ca-i). In addition, compared with control mice, both the *Nr2f1* and *Nr2f2* genes were almost completely deleted in the hippocampal primordium in mutant mice at E14.5 (Figure 1—figure supplement 1Cj-o). Since the LacZ expression serves as an indicator for the deletion of *Nr2f2* (Swindell et al., 2006; Tang et al., 2012), we performed immunofluorescence staining with antibodies against NR2F2 and LacZ on the sagittal sections of RX^Cre/+^; *Nr2f2^F/+^* and RX^Cre/+^; *Nr2f2^F/F^* mice at E11.5. NR2F2 was readily detected at the hippocampal primordium of the heterozygous mutant embryo at E11.5 (Figure 1—figure supplement 1Da, c, g); in contrast, the expression of *Nr2f2* was significantly reduced in the homozygous mutant (Figure 1—figure supplement 1Dd, f, j). In addition, compared with the heterozygous mutant embryo (Figure 1— figure supplement 1Db-c, h), the LacZ signals clearly increased in the hippocampal primordium of the homozygous mutant embryo at E11.5 (Figure 1—figure supplement 1De-f, k), suggesting that RX-Cre recombinase can efficiently excise the *Nr2f2* gene in the hippocampal primordium as early as E11.5. Intriguingly, we observed that the expression of *Nr2f1* increased in the caudal hippocampal primordium of the *Nr2f2* homozygous mutant embryo at E11.5 (Figure 1—figure supplement 1Di, l), indicating that similar to the observations in the early optic cup (Tang et al., 2010), *Nr2f1* and *Nr2f2* genes could be partially compensate with each other in the developing hippocampal primordium. All the data above show that RXCre recombinase could efficiently excise *Nr2f1* and/or *Nr2f2* in the early developing and postnatal hippocampus.

#### The *Nr2f2* gene is required for the appropriate morphogenesis of the ventral hippocampus but not of the dorsal hippocampus

Given that the *Nr2f2* gene is highly and specifically expressed in the postnatal ventral hippocampus and the CH of the hippocampal primordium (Figure 1 and Figure 1—figure supplement 1), we asked whether the *Nr2f2* gene is required for the appropriate morphogenesis of the hippocampus, particularly the ventral hippocampus. To answer this question, we conducted Nissl staining with samples from the *Nr2f2* single-gene (RX^Cre/+^; *Nr2f2^F/F^*) knockout mouse model. In coronal sections, compared with the control at 1M, the septal hippocampus was normal; unexpectedly, an ectopic CA-like region was observed medially in the temporal hippocampus in the *Nr2f2* mutant, where the prospective posterior part of the medial amygdaloid (MeP) nucleus was situated, indicated by the star (Figure 1Ba-f). The presence of the ectopic CA-like region in the ventral but not dorsal hippocampus of the mutant was further confirmed by the presence of the prospective MeP and amygdalohippocampal area (AHi) in sagittal sections, as indicated by the star (Figure 1Bg-l). Furthermore, immunofluorescence assays were performed to verify whether specific lineages of the hippocampus were altered with sagittal sections, in which the subregions of both the dorsal and ventral hippocampus were well displayed and distinguished. Ctip2 is a marker for CA1 pyramidal neurons and DG granule neurons, and HuB is a marker for CA3 pyramidal cells (Sugiyama, Osumi, & Katsuyama, 2014). The dorsal hippocampus appeared normal in both the control and mutant mice at 1M (Figure 1Ca-l). Nonetheless, compared with the observations in control mice, an ectopic HuB-positive CA3 pyramidal neuron lineage, indicated by the star, and a duplicated Ctip2-positive CA1 pyramidal neuron lineage, indicated by the arrowhead, were observed in the ventral hippocampal area in the mutant (Figure 1Ca-f, m-r), revealing that there were ectopic CA1 and CA3 lineages in the *Nr2f2* mutants.

Consistent with the previous report (Leid et al., 2004), the expression of Ctip2 was detected in the amygdala including the AHi and posteromedial cortical amygdaloid nucleus (PMCo); in addition, Ctip2 was also highly expressed in the dorsal part of the MeP (MePD) in the control (Figure 1Cb, n).

Intriguingly, compared with the controls at 1M, there were ectopic CA domains in the mutant ventral hippocampus with the expense of the Ctip2 positive AHi and MePD amygdaloid nuclei (Figure 1Ce, q), indicated by the arrows. Clearly, all the data above suggested that the *Nr2f2* gene is necessary to ensure the appropriate morphogenesis of the ventral hippocampus.

At early embryonic stages, *Nr2f2* was preferentially expressed in the CH (Figure 1—figure supplement 1Ab, d, 1Bb, e), the organizer of the hippocampus, and at postnatal 1-month-old (1M) stage, *Nr2f2* was also highly expressed in some amygdala nuclei such as the AHi and medial amygdaloid nucleus, which are adjacent to the ventral/temporal hippocampus (Figure 1Ae, h, n) (Tang et al., 2012). We would like to investigate the correlation of the CH and/or amygdala anlage with the duplicated ventral hippocampal domains in the *Nr2f2* mutant in detail in our future study. The observations above suggest that *Nr2f2* is not only specifically expressed in the ventral hippocampus but is also required for morphogenesis and probably the function of the ventral hippocampus. Since the ventral hippocampus participates in the regulation of emotion and stress, mutations in the *Nr2f2* gene lead to CHDs, and the formation of the ventral hippocampus is disrupted in *Nr2f2* mutant mice at 1M, we wondered whether CHD patients with *Nr2f2* mutations also exhibit symptoms associated with psychiatric disorders such as depression, anxiety, or schizophrenia.

#### The *Nr2f1* gene is required for the specification and differentiation of the dorsal CA1 identity

Next, we asked whether the deletion of the *Nr2f1* gene by RX-Cre also affected the development of the hippocampus. Consistent with the previous finding in *Emx1^Cre/+^*; *Nr2f1^F/F^*mutant mice (Flore et al., 2017), it was the septal/dorsal hippocampus, not the temporal/ventral hippocampus, that was specifically shrunken in both coronal (Figure 2Aa-f) and sagittal sections (Figure 2Ag-i) of RX^Cre/+^; *Nr2f1^F/F^* mutant mice. Then, we asked whether the loss of the *Nr2f1* gene caused abnormal specification and differentiation of hippocampal lineages. Compared with the control mice, *Nr2f1* mutant mice had fewer HuB-positive CA3 pyramidal neurons, as indicated by the star; intriguingly, Ctip2-positive CA1 pyramidal neurons failed to be detected, as indicated by the arrowhead, with Ctip2-positive DG granule neurons unaltered in the dorsal hippocampus (Figure 2Ba-l). The loss of the dorsal CA1 pyramidal neuron identity in mutant mice was further confirmed by Wfs1, another dorsal CA1 pyramidal neuron-specific marker (Takeda et al., 2001) (Figure 2Ca-r). Nonetheless, the HuB-positive and Ctip2-positive lineages were comparable in the ventral hippocampus between the control and mutant mice (Figure 2Ba-f, m-r), even though the low expression of NR2F1 was detected there. Indeed, *Nr2f1* is not only expressed at the highest level in the dorsal CA1 but is also required for the specification and differentiation of dorsal CA1 pyramidal neurons, among which place cells are essential for learning and memory (O’Keefe & Conway, 1978; O’Keefe & Dostrovsky, 1971).

Given that dysplasia of the dorsal hippocampus was generated in both the *Emx1^Cre^* and RX^Cre^ models (Flore et al., 2017) (Figure 2Aa-l), we asked whether the development of dorsal CA1 pyramidal neurons was also abolished in *Emx1^Cre/+^*^;^ *Nr2f1^F/F^* mutant mice. To answer this question, immunofluorescence staining was conducted first. Compared with that in the control mice, the proportions of either the Wfs1- or Ctip2-positive CA1 domain were reduced in the mutant mice at 3M (Figure 2—figure supplement 1Aa-h), indicating that the differentiation of the dorsal CA1 pyramidal neurons was also compromised in the *Emx1^Cre^* model, although it was less severe than that in the RX^Cre^ model. To make our findings more consistent with previous studies, we further conducted experiments with the *Emx1^Cre^* model. Afterward, to investigate the fine structure of the dorsal CA1 pyramidal neurons, Golgi staining was performed. Compared with those of control mice, the numbers of secondary dendrites and branch points of both the apical and basal dendrites were significantly reduced in the dorsal CA1 pyramidal neurons of mutant mice at 3M (Figure 2—figure supplement 1Ba-e). Then, the dorsal hippocampus-related spatial learning and memory behavior test, the Morris water maze, was performed (Vorhees & Williams, 2006). Consistent with a previous report (Flore et al., 2017), spatial learning and memory function was significantly impaired in adult *Emx1^Cre/+^*; *Nr2f1^F/F^* mice, compared with the control mice (Figure 2—figure supplement 1C). The data above suggest that *Nr2f1* is vital for the morphogenesis, lineage specification, and spatial learning and memory of the dorsal hippocampus, and particularly, the compromised dorsal CA1 lineage could contribute to the phenotypes associated with neurodevelopmental disorders, including ID or ASD.

**Figure 2.**
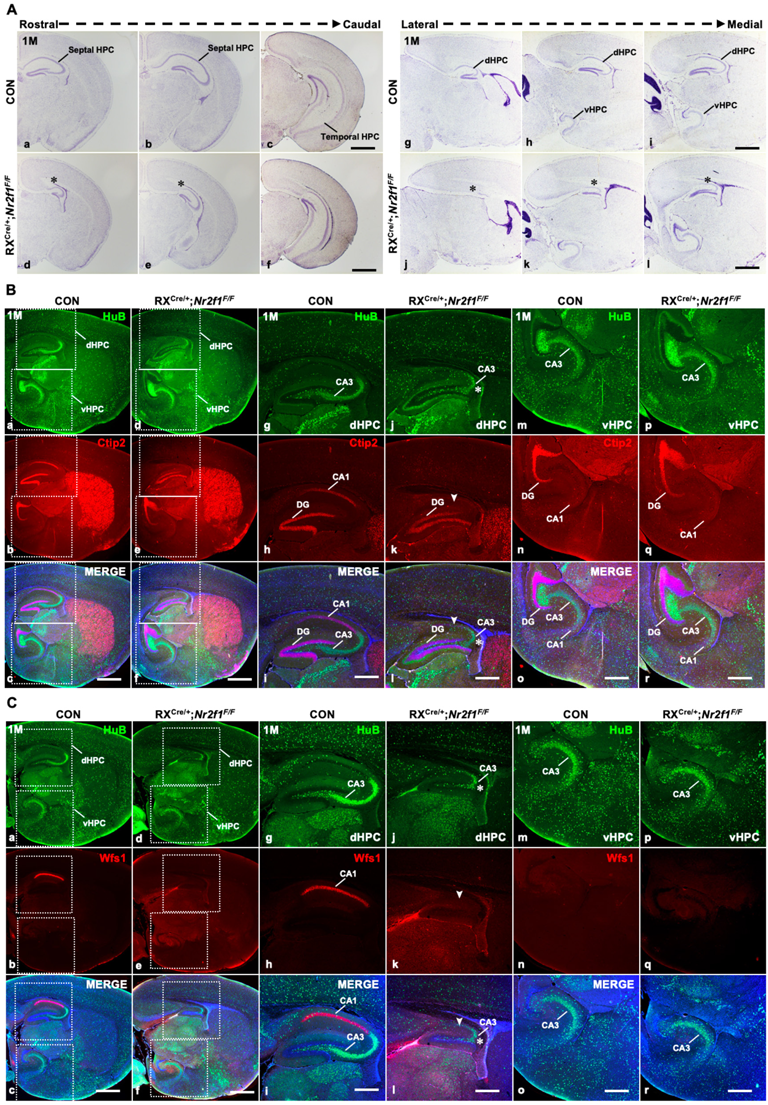
The specification and differentiation of the dorsal CA1 lineage failed with the dysplastic dorsal hippocampus in RX^Cre/+^; *Nr2f1^F/F^* mutant mice. A, In coronal sections along the rostrocaudal axis (**a-f**) and sagittal sections along the lateral-medial axis (**g-l**) of the hippocampus, compared with that of control mice (**a-c**, **g-i**), the dorsal hippocampus was shrunken, indicated by the star, in *Nr2f1* gene mutant (RX^Cre/+^; *Nr2f1^F/F^)* mice at 1M (**d-f**, **j-l**). **B,** The expression of HuB and Ctip2 in the corresponding inserted area in **a-f** under a high magnification objective lens at 1M (**g-r**); compared with that of control mice (**a-c**, **g-i**, **m-o**), the HuB-positive CA3 domain was reduced in the dorsal hippocampus, especially the Ctip2-positive dorsal CA1, which was barely detected in *Nr2f1* mutant mice at 1M (**d-f**, **j-l**), while their expression in the ventral hippocampus was comparable between the controls and mutants (**d-f**, **p-r**). **C,** The expression of HuB and Wfs1 in the corresponding inserted area in **a-f** under a high magnification objective lens at 1M (**g-r**); the expression of HuB and the dCA1 marker Wfs1 in the control (**a-c**, **g-i**, **m-o**) and *Nr2f2* mutant mice (**d-f**, **j-l**, **p-r**) at 1M. Wfs1-positive dorsal CA1 could not be detected in *Nr2f1* mutant mice at 1M, as indicated by the arrowhead. dHPC, dorsal hippocampus; HPC, hippocampus; vHPC, ventral hippocampus. Scale bars, **Aa-l**, **Ba-f**, **Ca-f,** 200 μm; **Bg-r**, **Cg-r,** 100 μm.

#### The *Nr2f1* and *Nr2f2* genes coordinate to ensure the genesis of the hippocampus

Given that the loss of either *Nr2f1* or *Nr2f2* leads to dysplasia of the dorsal or ventral hippocampus, respectively, we asked whether these genes compensate for each other to regulate the morphogenesis of the hippocampus. To answer this question, the RX^Cre/+^; *Nr2f1^F/F^*; *Nr2f2^F/F^*double-mutant mouse was generated, and a few homozygous double-gene mutant mice survived for approximately 3 weeks (3W). Nonetheless, the reason for the lethality of the double-mutant mice is still unknown. Nissl staining data showed that compared with that of control mice, the septal hippocampus was severely shrunken, as indicated by the star, and the temporal hippocampus was barely observed in the double-mutant mouse brains (Figure 3Aa-h). Unexpectedly, an ectopic nucleus was observed in the region of the prospective temporal hippocampus, indicated by the arrowhead, in the double-mutant mice (Figure 3Ag-h). In addition, compared with those of controls, the regions with HuB-positive CA3 pyramidal neurons and Ctip2-positive or Prox1-positive DG granule neurons were diminished in the double mutants; in particular, Ctip2-positive dorsal CA1 pyramidal neurons could not be detected in the double mutants (Figure 3Ba-l). Furthermore, compared with the domains of controls, no HuB-positive, Ctip2-positive, or Prox1-positive domains could be detected in the prospective temporal hippocampus in the double mutants (Figure 3Ca-l). The results above suggest that the *Nr2f1* and *Nr2f2* genes coordinate with each other to mediate the appropriate morphogenesis of the entire hippocampus.

**Figure 3.**
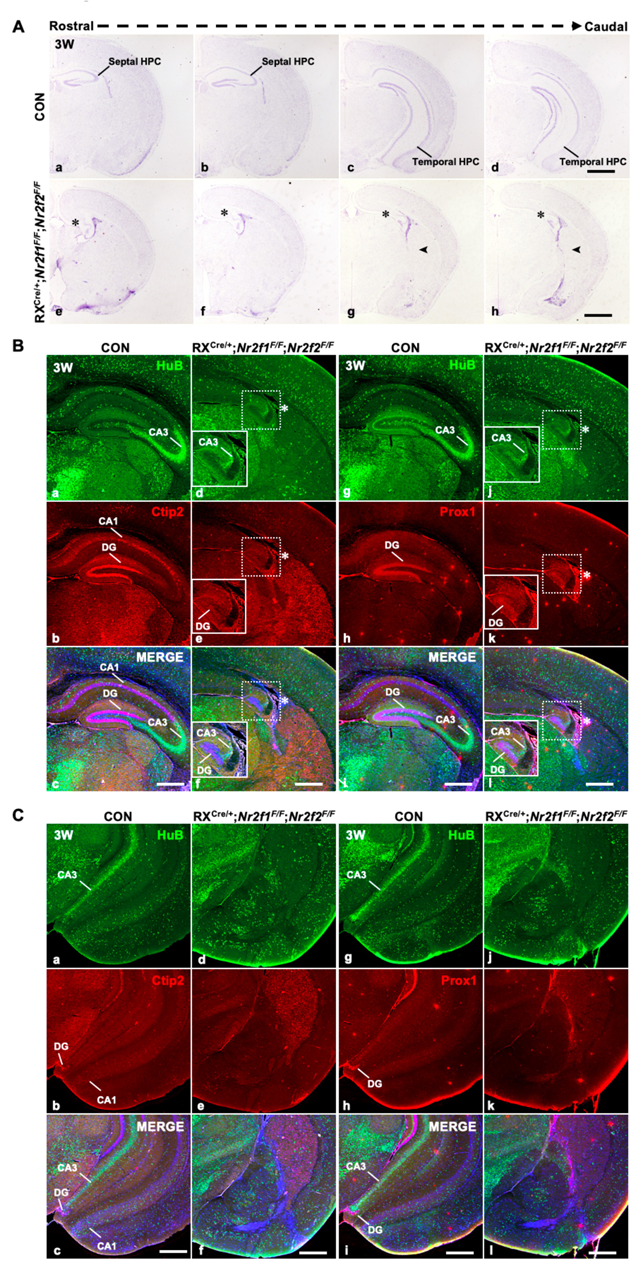
Defects in the hippocampus in RX^Cre/+^; *Nr2f1^F/F^*; *Nr2f2^F/F^*double*-*gene mutant mice. **A**, In coronal sections along the rostrocaudal axis, compared with control mice (**a-d**), the hippocampus was atrophic in RX^Cre/+^; *Nr2f1^F/F^; Nr2f2^F/F^* double mutant mice, indicated by the star, and an ectopic unknown nucleus was observed in the caudal plates, indicated by the arrowhead (**e-h**). **B,** Compared with that of control mice (**a-c**, **g-i**), the expression of HuB, Ctip2, and Prox1 was decreased in the hippocampus of *Nr2f1/2* double-gene mutant mice at 3 weeks postnatal (3W) (**d-f**, **j-l**). **C,** Compared with that of control mice (**a-c**, **g-i**), the expression of HuB could not be detected in the presumptive CA3 domain, and the expression of Ctip2 or Prox1 could not be detected in the presumptive DG domain of the prospective ventral hippocampus of RX^Cre/+^; *Nr2f1^F/F^*; *Nr2f2^F/F^* double mutant mice. Scale bars, **Aa-h**, 200 μm; **Ba-l**, **Ca-l,** 100 μm.

#### *Nr2f* genes and adult neurogenesis in the hippocampus

Given that the *Nr2f1* or *Nr2f2* gene was highly expressed in the dorsal or ventral DG, respectively (Figure 1A), and that RX-Cre recombinase could efficiently delete either gene in the DG (Figure 1—figure supplement 1Ca-i), we asked whether the loss of the *Nr2f1* or/and *-Nr2f2* gene in the DG would affect hippocampal adult neurogenesis. To answer this question, the ventral DG, dorsal DG, and septal DG were chosen to perform immunofluorescence assays in the *Nr2f2* mutant, *Nr2f1* mutant, and double mutant models, respectively. Adult NSCs in the subgranular zone (SGZ) of the DG express both GFAP and Nestin, and newborn granule neurons express Dcx (Gao, Arlotta, Macklis, & Chen, 2007). The numbers of NSCs and newborn neurons in the SGZ of the DG were comparable between control mice and either the *Nr2f2* or *Nr2f1* mutant mice (Figure 3—figure supplement 1Aa-p, Ba-h); nonetheless, compared with those of control mice, the numbers of the NSCs and newborn granule neurons in the SGZ of the DG were reduced in the double mutants (Figure 3—figure supplement 1Aq-x, Bi-l), and the reduction in both lineages was significant (Figure 3—figure supplement 1Ca, b). The data above suggest that *Nr2f* genes may coordinate with each other to execute essential functions for appropriate hippocampal adult neurogenesis in the DG.

### Hippocampal trisynaptic connectivity was impaired in postnatal *Nr2f2* single-, *Nr2f1* single-, and double-mutant mice at about 1M

Given that dysplasia of the hippocampus was observed in all three mouse models, we asked whether the connectivity of the hippocampal trisynaptic circuit associated with the DG, CA3, and CA1 regions (Amaral, 1993) was normal in these models. To answer this question, the components of the trisynaptic circuit were characterized in the ventral hippocampus of *Nr2f2* mutants, the dorsal hippocampus of *Nr2f1* mutants, and the septal hippocampus of double mutants. Calretinin is a marker of mossy cells, Calbindin is a marker of mossy fibers, and SMI312 is a marker of Schafer collaterals (Flore et al., 2017). Compared with those of controls (Figure 4Aa, b, e, f, i, and j), the numbers of Calretinin-positive mossy cells were reduced, Calbindin-positive mossy fibers were longer but thinner, and SMI312-positive Schafer collaterals were thinner and discontinued in the ventral hippocampus of *Nr2f2* mutants at 1M (Figure 4Ac, d, g, h, k, and l). In addition, similar to the previous report (Flore et al., 2017), the numbers of Calretinin-positive mossy cells were decreased, Calbindin-positive mossy fibers were shorter and thinner, and SMI312-positive Schafer collaterals were barely detected in the dorsal hippocampus of *Nr2f1* mutants (Figure 4Ac, d, g, h, k, and l), compared with those of controls at 1M (Figure 4Ba, b, e, f, i, and j). Moreover, compared with those of control mice (Figure 4Ca, b, e, f, i, and j), the numbers of Calretinin-positive mossy cells were reduced; both Calbindin-positive mossy fibers and SMI312-positive Schafer collaterals were barely detected in the prospective septal hippocampus of the double mutants at 3W (Figure 4Cc, d, g, h, k, and l). Clearly, without both *Nr2f* genes, the connectivity of the hippocampal trisynaptic circuit was abolished more severely. The observations above revealed that the formation of the trisynaptic circuit, which is one of the fundamental characteristics of hippocampal neurophysiology (Basu & Siegelbaum, 2015), was abnormal in all three mouse models, indicating that both the morphology and functions of the hippocampus are most likely compromised in the loss of the *Nr2f1* and/or *Nr2f2* gene.

**Figure 4.**
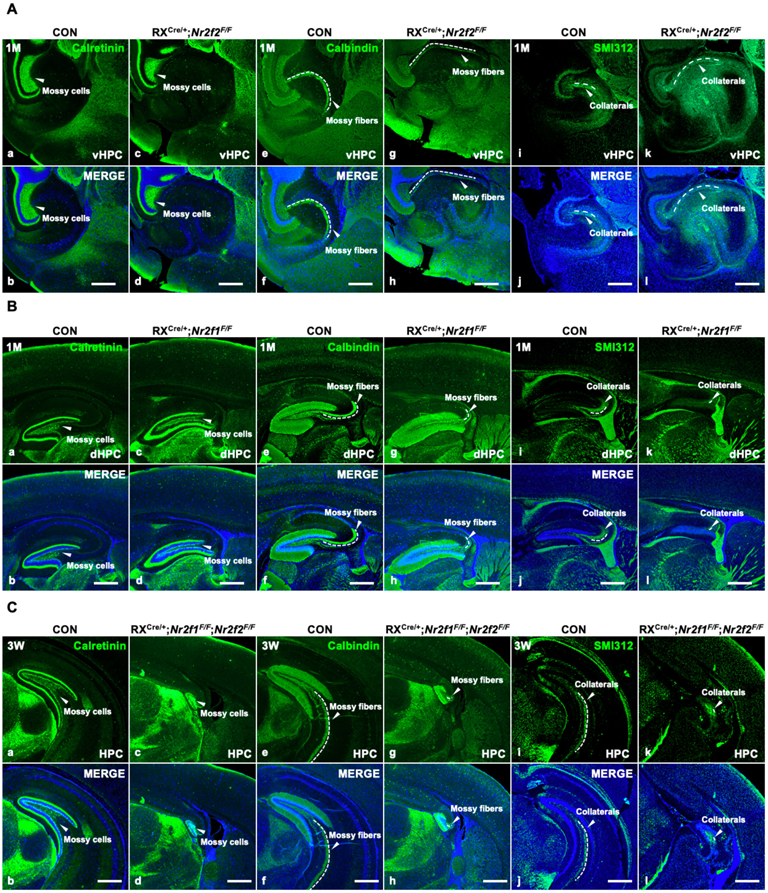
The impairment of hippocampal trisynaptic connectivity in *Nr2f2* single-gene, *Nr2f1* single-gene, and *Nr2f1/2* double-gene mutant mice. A, The expression of Calretinin, Calbindin, and SMI312 in the ventral hippocampus of the control (**a-b**, **e-f**, **i-j**) and *Nr2f2* single-gene mutant mice (**c-d**, **g-h**, **k-l**). **B,** The expression of Calretinin, Calbindin, and SMI312 in the dorsal hippocampus of the control (**a-b**, **e-f**, **i-j**) and *Nr2f1* single-gene mutant mice (**c-d**, **g-h**, **k-l**). **C,** The expression of Calretinin, Calbindin, and SMI312 in the hippocampus of the control (**a-b**, **e-f**, **i-j**) and *Nr2f1*/*2* double-gene mutant mice (**c-d**, **g-h**, **k-l**). dHPC, dorsal hippocampus; HPC, hippocampus; vHPC, ventral hippocampus. Scale bars, **Aa-l**, **Ba-l**, **Ca-l**, 100 μm.

### The expression of several essential regulatory genes associated with early hippocampal development was abnormal in double mutants

Given that the hippocampus was almost completely diminished in double mutants, we asked how *Nr2f* genes participated in the regulation of the early morphogenesis of the hippocampus. To answer this question, total RNA isolated from the whole telencephalons of control (n=5) and double-mutant-(n=3) embryos at E11.5 was used to generate cDNA, and then real-time quantitative PCR (RT-qPCR) assays were performed. As expected, compared with that of control mice, the expression of *Nr2f1* and *Nr2f2* was reduced significantly in the double mutant mice (Figure 5A). Then, we mainly focused on the intrinsic regulatory networks by analyzing the expression profiles of two groups of transcription factor genes. The *Foxg1*, *Gli3*, *Lhx2*, *Otx1, Otx2*, and *Pax6* genes, which are highly related to the early patterning of the dorsal telencephalon (Hebert & Fishell, 2008), were in the first group; *Axin2*, *Emx1*, *Emx2*, *Lef1*, *Lhx5*, and *Tcf4* genes, which are associated with early hippocampal development (Galceran et al., 2000; Moore & Iulianella, 2021; Tole et al., 2000; Yoshida et al., 1997; Zhao et al., 1999), were in the other group. The expression of the *Foxg1*, *Gli3*, *Lhx2*, *Otx1, Otx2*, and *Pax6* genes was comparable between the controls and double mutants (Figure 5A), indicating that the early patterning of the dorsal telencephalon is largely unaltered. Compared with that of control mice, the expression of the *Axin2*, *Emx2*, *Lef1*, and *Tcf4* genes was normal in the double mutants; interestingly, the expression of the *Emx1* and *Lhx5* transcripts was decreased significantly in the double mutants at E11.5 compared to that in control mice (Figure 5A). Consistent with the downregulated expression of *Lhx5* transcripts in the double mutant, the expression of the Lhx5 protein was reduced in the CH in the double mutants at E11.5; moreover, the number of Lhx5-positive Cajal-Retzius cells decreased in the double mutant embryos at E11.5, E13.5 and E14.5 (Figure 5Ba-d, a’-d’, a’’-d’’, i-l, i’-l’, q-t, q’-t’). The expression of *Lhx2* was expanded ventrally into the choroid plexus in the *Lhx5* null mutant mice (Zhao et al., 1999), indicating that *Lhx5* could inhibit *Lhx2* expression locally. Consistent with RT-qPCR data, the expression of Lhx2 was comparable between the control and double-mutant mice at E11.5 (Figure 5Be-h, e’-h’). Interestingly, the expression of the Lhx2 protein was increased in the hippocampal primordium in the *Nr2f* double-mutant mice at E13.5 and E14.5 (Figure 5Bm-p, m’-p’, u-x, u’-x’). The upregulation of Lhx2 expression is most likely associated with the reduced expression of the *Lhx5* gene.

**Figure 5.**
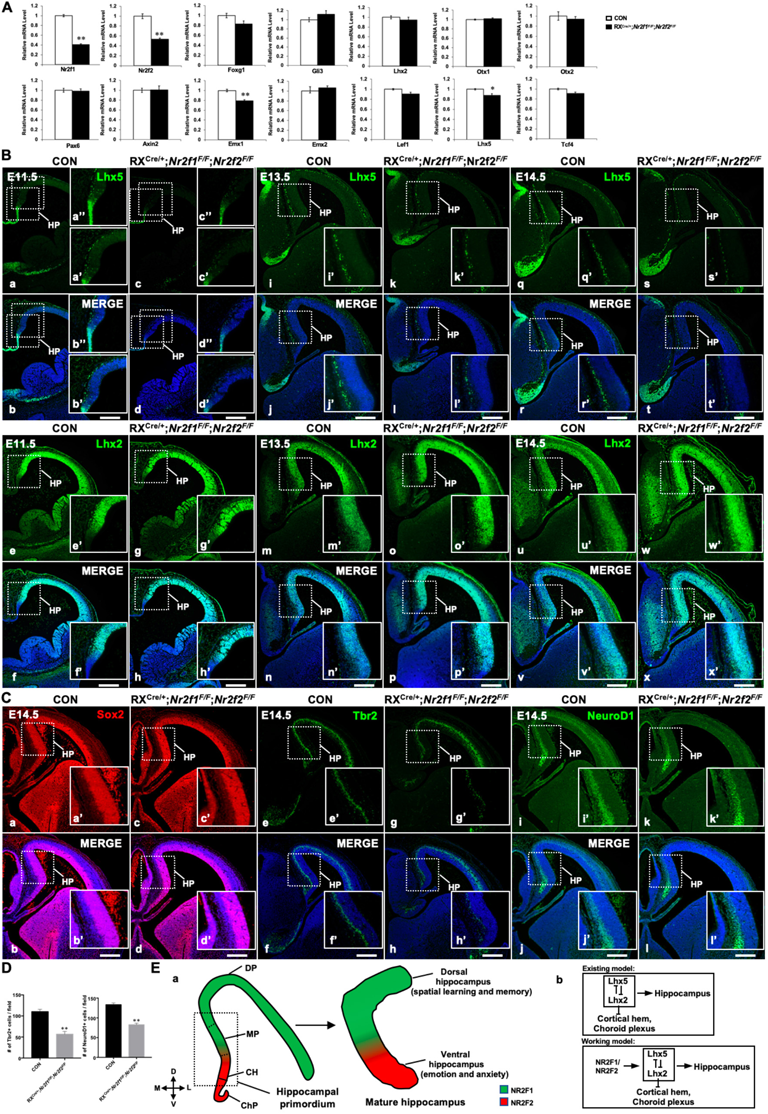
*Nr2f* genes regulate the expression of key genes associated with early hippocampal development. **A**, The expression profiles of genes involved in hippocampal development in control and the double mutant mice at E11.5. **B,** Compared with that of control mice (**a-b**, **a’-b’, a’’-b’’, i-j, i’-j’, q-r, q’-r’**), the expression of Lhx5 was reduced in double-mutant mice at E11.5 (**c-d**, **c’-d’, c’’-d’’**), E13.5 (**k**-**l**, **k’**-**l’**) and E14.5 (**s**-**t**, **s’**-**t’**); the expression of Lhx2 was comparable between the control and double-mutant mice at E11.5 (**e**-**h**, **e’**-**h’**); and compared with that of control mice (**m-n**, **m’**-**n’**, **u**-**v**, **u’-v’**), the expression of Lhx2 was increased in double-mutant mice at E13.5 (**o**-**p**, **o’**-**p’**) and E14.5 (**w-x**, **w’-x’**). **C,** Compared with that of control mice (**a-b**, **a’-b’**), the expression of Sox2 was normal in double-mutant mice at E14.5 (**c-d**, **c’-d’**); compared with that of control mice (**e-f**, **e’-f’**), the expression of Tbr2 was decreased in *Nr2f* mutant mice at E14.5 (**g-h**, **g’-h’**); compared with that of control mice (**i-j**, **i’-j’**), the expression of NeuroD1 was reduced in double-mutant mice at E14.5 (**k-l**, **k’-l’**). **D,** Quantitative analysis of Tbr2-positive cells and NeuroD1-positive cells in **Ce’-h’** and **Ci’-l’**. **E,** In the hippocampal primordium of the early embryo, *Nr2f1* is expressed dorsally in the MP, and *Nr2f2* is expressed ventrally in the CH. In the mature hippocampus, the expression of *Nr2f1* is higher in the dorsal hippocampus, which is related to spatial learning and memory, and the expression of *Nr2f2* is mainly in the ventral hippocampus, which is associated with emotion and anxiety (**a**). Our findings support a novel molecular mechanism by which *Nr2f1* and *Nr2f2* may cooperate to ensure the appropriate morphogenesis and functions of the hippocampus by modulating the *Lhx5*-*Lhx2* axis (**b**). Data are expressed as the mean ± SEM. *Student’s t test*, \*\**P*<0.01. CH, cortical hem; ChP, choroid plexus; DP, dorsal pallium; HP, hippocampal primordium; MP, medial pallium. Scale bars, **Ba-x, Ca-l**, 200 μm.

Next, we asked whether neural precursor cells (NPCs), intermediate progenitor cells (IPCs), or newborn neurons were affected in the early development of the hippocampus in double-mutant mice. Sox2 is a marker for NPCs, Tbr2 is a marker for IPCs, and NeuroD1 is a marker for newborn neurons (Yu, Marchetto, & Gage, 2014). The expression of Sox2 in the hippocampal regions was comparable between the control and double-mutant mice at E14.5 (Figure 5Ca-d, a’-d’), indicating that the generation of NPCs was normal. Nevertheless, compared with the control embryos, the numbers of Tbr2-positive IPCs and NeuroD1-positive newborn neurons were reduced in the double-mutant embryos (Figure 5Ce-l, e’-l’), and the reduction was significant (Figure 5D). Our observations were consistent with previous findings in *Lhx5* null mutant mice that the specification of the hippocampal NPCs was normal, but the later differentiation event was abolished (Zhao et al., 1999). All the data above suggest that *Nr2f* genes may cooperate to ensure the early morphogenesis of the hippocampus by regulating the appropriate expression levels of *Lhx5* and *Lhx2* genes. Nevertheless, we could not exclude other possibilities that *Nr2f* genes could also participate in the modulation of hippocampal development through *Emx1* or other genes.

## Discussion

In our present study, we observed dorsal-high NR2F1 and ventral-high NR2F2 expression profiles in the postnatal hippocampus. The deletion of the *Nr2f2* gene led to duplicated CA1 and CA3 domains of the ventral hippocampus. The loss of *Nr2f1* resulted in the failed specification and differentiation of the dorsal CA1 pyramidal neuron lineage with a diminished dorsal hippocampus. Furthermore, the deficiency of both *Nr2f* genes caused atrophy of almost the entire hippocampus, accompanied by compromised generation of the CA1, CA3, and DG identities. In addition, the dorsal trisynaptic components, ventral trisynaptic components, or entire trisynaptic components were abolished in the corresponding *Nr2f1* gene mutant, *Nr2f2* gene mutant, or *Nr2f1*/*2* double-gene mutant mice. Moreover, *Nr2f* genes may cooperate to ensure the appropriate morphogenesis and function of the hippocampus by regulating the *Lhx5-Lhx2* axis.

### 1. *Nr2f2* governs the distinct characteristics of the ventral hippocampus

Sixty years ago, the pioneering work of Milner and her colleagues discovered the essential role of the hippocampus in declarative memory (Penfield & Milner, 1958; Scoville & Milner, 1957). Recently, accumulating evidence has supported the Moser theory that the hippocampus is a heterogeneous structure with distinct characteristics of gene expression, connectivity, and function along its dorsoventral axis (Bast, 2007; Fanselow & Dong, 2010; Moser & Moser, 1998; Strange et al., 2014). The dorsal hippocampus marked in blue, in which gene expression is similar to the neocortex, serves the “cold” cognitive function associated with declarative memory and spatial navigation, and the ventral hippocampus marked in red, in which gene expression is close to the hypothalamus and amygdala, corresponds to the “hot” affective states related to emotion and anxiety (Figure 5—figure supplement 1). The ventral hippocampus generates direct connectivity with the amygdala, hypothalamus, medial prefrontal cortex (mPFC), and olfactory bulb (Cenquizca & Swanson, 2007; Hoover & Vertes, 2007; Kishi et al., 2000; Pitkanen et al., 2000; Roberts et al., 2007). Nonetheless, thus far, the molecular and cellular mechanism of how the morphogenesis, connectivity, and function of the ventral hippocampus is achieved has been largely unclear.

*Nr2f2*, a nuclear receptor gene associated with heart disease(Al Turki et al., 2014) (High et al., 2016), was highly and exclusively expressed in the ventral hippocampus in 1-month-old mice and was expressed ventrally in the CH of the hippocampal primordium in mouse embryos (Figure 1, Figure 1—figure supplement 1, Figure 5Ea), indicating that the *Nr2f2* gene may participate in the regulation of the development and function of the ventral hippocampus. First, deficiency of the *Nr2f2* gene led to the duplication of the CA1 and CA3 domains of the ventral hippocampus but not the dorsal hippocampus, which was confirmed both morphologically and molecularly (Figure 1). Second, the formation of the trisynaptic circuit was specifically abolished in the ventral hippocampus of *Nr2f2* mutants (Figure 4), indicating that the intrahippocampal circuit, information transfer, and function of the ventral hippocampus could be compromised. Third, the ventral hippocampus generates neural circuits with the mPFC, amygdala, nucleus accumbens, and hypothalamus, which are associated with anxiety/behavioral inhibition, fear processing, pleasure/reward seeking, and the neuroendocrine system, respectively (Anacker & Hen, 2017; Baik, 2020; Bryant & Barker, 2020; Cenquizca & Swanson, 2007; Herman et al., 2016; Kishi et al., 2000; O’Leary & Cryan, 2014; Pitkanen et al., 2000). These ventral hippocampal projections may be important for processing information related to emotion and anxiety. Intriguingly, our previous studies revealed that *Nr2f2* is required for the development of the hypothalamus, amygdala, and olfactory bulb (S. Feng et al., 2017; Tang et al., 2012; X. Zhou et al., 2015), all of which generate functional neural circuits with the ventral hippocampus (Fanselow & Dong, 2010). Particularly, both the hypothalamus and amygdala are also diminished in RX^Cre/+^; *Nr2f2^F/F^* mutant mice (S. Feng et al., 2017; Tang et al., 2012), indicating that their interconnectivities with the ventral hippocampus are abnormal. Thus, all the findings above suggest that *Nr2f2* is a novel and essential intrinsic regulator that controls the morphogenesis, connectivity, and function of the ventral hippocampus.

Given that mutations of *Nr2f2* are highly associated with CHDs (Al Turki et al., 2014), the expression of *Nr2f2* is also confined to the ventral hippocampus in human embryos (Alzu’bi et al., 2017), and *Nr2f2* gene is required for the distinct characteristics of the ventral hippocampus in mouse (Figure 1, Figure 4), we wondered whether CHD patients carrying mutations of *Nr2f2* also display symptoms of psychiatric disorders, such as depression, anxiety, or schizophrenia, related to the ventral hippocampus. In our future study, we would like to generate hippocampus-specific or hippocampal subdomain-specific conditional knockout models to dissect distinct roles of the *Nr2f2* gene in the hippocampus, particularly in the ventral hippocampus, in detail.

### 2. The *Nr2f1* gene is required for the specification and differentiation of dorsal CA1 pyramidal neurons

The expression of *Nr2f1*, another orphan nuclear receptor gene associated with neurodevelopmental disorders (Bertacchi et al., 2020; Bosch et al., 2014; Contesse et al., 2019), is high in the dorsal MP of the hippocampal primordium and is higher in the dorsal hippocampus than in the ventral hippocampus (Figure 1, Figure 1—figure supplement 1, Figure 5Ea) (Flore et al., 2017). Consistent with previous observations in *Emx1^Cre/+^*; *Nr2f1^F/F^* mutant mice (Flore et al., 2017), the dorsal hippocampus but not the ventral hippocampus was specifically shrunken in RX^Cre/+^; *Nr2f1^F/F^* mice (Figure 2, Figure 5—figure supplement 1). *Nr2f1* is expressed at the highest level in dorsal CA1 pyramidal neurons (Figure 1), indicating that the *Nr2f1* gene may play a role in the specification and differentiation of dorsal CA1 pyramidal neurons. As expected, the expression of Ctip2 and Wfs1, two markers for dorsal CA1 pyramidal neurons, could not be detected in the prospective dorsal CA1 domain in RX^Cre/+^; *Nr2f1^F/F^*mutant mice (Figure 2); furthermore, *Emx1^Cre/+^*; *Nr2f1^F/F^* mutant mice partially phenocopied the compromised development of the dorsal CA1 lineage (Figure 2—figure supplement 1). It seems that the spatiotemporal activity of RX-Cre recombinase is better or broader than that of the Emx1-Cre recombinase during the critical period of the specification of the dorsal CA1 pyramidal neuron identity. In addition, the Golgi staining assay revealed that the development of the dendrites of the dorsal CA1 pyramidal neurons was abnormal (Figure 2—figure supplement 1). All the observations above indicate that the *Nr2f1* gene is not only necessary for the morphogenesis of the dorsal hippocampus but is also required for the specification and differentiation of the dorsal CA1 pyramidal neurons, among which there are place cells. The identification of place cells fifty years ago was one of the most important breakthroughs in understanding the role of the hippocampus in memory (O’Keefe & Dostrovsky, 1971). Except for spatial information, place cells in the dorsal CA1 may also encode nonspatial representations, such as time (Eichenbaum, 2017; Lisman et al., 2017). Notably, 95% of patients carrying *Nr2f1* mutations are associated with ID. Here, our observations support the notion that the *Nr2f1* gene is a novel intrinsic regulator that specifies the dorsal CA1 pyramidal cell identity, which will benefit the understanding of both neurophysiological functions of the hippocampus and the etiology of NDD including ID and ASD.

### 3. *Nr2f1* and *Nr2f2* cooperate to ensure the appropriate morphogenesis of the hippocampus by regulating the *Lhx5*-*Lhx2* axis in mice

In wild-type mice, *Nr2f1* and *Nr2f2* genes generated complementary expression profiles in the embryonic hippocampal primordium with *Nr2f1* in the dorsal MP, marked in green; *Nr2f2* in the ventral CH, marked in red (Figure 5Ea, Figure 1—figure supplement 1); and in the postnatal hippocampus with high-NR2F1 expression in the dorsal, marked in green; and high-NR2F2 expression in the ventral, marked in red (Figure 5Ea, Figure 1). These findings indicated that *Nr2f* genes may coordinate to regulate hippocampal development. Indeed, as discussed above, the loss of either *Nr2f1* or *Nr2f2* only leads to dysplasia of the dorsal hippocampus (Flore et al., 2017) (Figure 2, Figure 2—figure supplement 1) or ventral hippocampus (Figure 1), respectively; intriguingly, while both genes are efficiently excised by RX-Cre in the hippocampal primordium (Figure 1—figure supplement 1), more severely shrunken hippocampi developed in the 3-week-old double knockout mice (Figure 3). The dosage-dependent severity of hippocampal abnormalities suggested that two nuclear receptor genes, *Nr2f1* and *Nr2f2* could cooperate with each other to execute an essential and intrinsic function in the development of the hippocampus.

It is known that both extrinsic signals and intrinsic factors participate in the regulation of the early development of the hippocampus. Notably, mutations of *Wnt3a* and *Lef1* eliminate the entire hippocampus (Galceran et al., 2000; S. M. Lee et al., 2000). Given that the expression of *Axin2*, *Lef1* and *Tcf4*, three Wnt-responsive transcription factor genes, was not altered in the *Nr2f* double mutant (Figure 5A), it is unlikely that abnormal Wnt signaling is the cause of the compromised hippocampus. *Lhx5* is specifically expressed in the hippocampal primordium and is required for the morphogenesis of the hippocampus (Zhao et al., 1999). *Lhx2* is necessary for hippocampal development by repressing cortical hem fate (Mangale et al., 2008; Monuki et al., 2001). Agenesis of the hippocampus is observed in either *Lhx5* or *Lhx2* null mutant mice, and these genes particularly repress each other (Hebert & Fishell, 2008; Mangale et al., 2008; Roy et al., 2014; Zhao et al., 1999), indicating that the proper expression levels of *Lhx5* and *Lhx2* genes are critical to maintain the appropriate development of the hippocampus (Figure 5Eb). The transcriptional and protein expression levels of *Lhx5* but not *Lhx2* were first reduced in the hippocampal primordium of *Nr2f* double-mutant mice at E11.5; later, enhanced expression of the Lhx2 protein was detected in the hippocampal primordium of double-mutant mice at E13.5 and E14.5 (Figure 5A-B). Moreover, the number of Lhx5-positive Cajal-Retzius cells was clearly reduced in the double mutant embryos at E11.5, E13.5 and E14.5; consistent with the observations in the *Lhx5* null mutant (Li et al., 2021; Miquelajáuregui et al., 2010), the generation of Sox2-positive hippocampal NPCs was not affected, but the development of Tbr2-positive IPCs and NeuroD1-positive newborn neurons was abnormal in *Nr2f* double-mutant mice (Figure 5). Thus, our findings reveal a novel intrinsic regulatory mechanism that *Nr2f1* and *Nr2f2*, two disease-associated nuclear receptor genes, may cooperate with each other to ensure proper hippocampal morphogenesis by regulating the *Lhx5-Lhx2* axis. Intriguingly, compared with the adult *Emx1^Cre/+^*; *Nr2f1^F/F^* mutant mice, the hippocampus was much smaller in the adult *Emx1^Cre/+^*; *Nr2f1^F/F^*; *Nr2f2^F/F^* double-gene mutant mice; nevertheless, both the dorsal and ventral hippocampus were readily detected in double-gene mutants with *Emx1^Cre^* (our unpublished observations). In addition, the discrepancy between the shrunken dorsal hippocampus associated with the loss of *Nr2f1* and the duplicated CA domains of the ventral hippocampus associated with the deficiency of *Nr2f2* suggested that the regulatory network related to *Nr2f* genes during the early morphogenesis of the hippocampus could be much more complicated than suspected and should be investigated in our future study.

### 4. *Nr2f* genes are imperative for the formation of the trisynaptic circuit

The hippocampus and entorhinal cortex (EC) are interconnected through various neural circuits to mediate the flow of the information associated with declarative memory (Basu & Siegelbaum, 2015). Both direct and indirect glutamatergic circuits are involved in the relay of information from the EC to the hippocampal CA1, and the trisynaptic pathway is the most well-characterized indirect circuit. The EC sends sensory signals from association cortices via the perforant path to the DG, then the DG granule cells send excitatory mossy fiber projections to CA3 pyramidal neurons, and CA3 pyramidal neurons project to CA1 via the Schaffer collaterals (H. Lee, GoodSmith, & Knierim, 2020). Consistent with the high expression of NR2F1 in the dorsal hippocampus and NR2F2 in the ventral hippocampus (Figure 1), the dorsal trisynaptic circuit is specifically damaged in *Nr2f1* mutants, as is the ventral trisynaptic circuit in *Nr2f2* mutant mice. Moreover, the hippocampal trisynaptic circuit was almost completely absent in the double mutants (Figure 4, and our unpublished observations). The information transfer associated with the trisynaptic circuits should be abolished particularly in the dorsal and/or the ventral hippocampus in the above corresponding genetic mouse models. Interestingly, *Nr2f1* is also required to specify the medial EC cell fate (J. Feng et al., 2021). Therefore, the impaired formation and function of trisynaptic circuits could be caused by the abnormal development of CA1, CA3, DG or EC lineages. Nonetheless, given that newborn granule neurons are continuously generated in the adult DG to integrate into the existing neural circuits essential for declarative memory (Toda, Parylak, Linker, & Gage, 2019; Tuncdemir, Lacefield, & Hen, 2019) and that hippocampal adult neurogenesis was severely compromised in the *Nr2f* double mutant (Figure 3—figure supplement 1), we could not exclude the possibility that impaired adult neurogenesis may also contribute to the malformation and impaired function of the trisynaptic pathway in double mutants. We would like to investigate what sort of synaptic circuitry is compromised either physiologically or morphologically in the trisynaptic circuit of individual animal model in detail in future studies.

The hippocampus is heterogeneous along its dorsoventral axis, and either the dorsal or ventral hippocampus generates unique and distinguishable characteristics of gene expression and connectivity, which enable the hippocampus to execute an integrative function from the encoding and retrieval of certain declarative memory to adaptive behaviors. Lesions of the dorsal hippocampus, which are essential for the cognitive process of learning and memory, lead to amnesia and ID; while damage to the ventral hippocampus, which is central for emotion and affection, is highly associated with psychiatric disorders including depression, anxiety, and schizophrenia. Our findings in this study reveal novel intrinsic mechanisms by which two nuclear receptor genes, *Nr2f1* and *Nr2f2*, which are associated with NDD or CHD, converge to govern the differentiation and integration of the hippocampus along the dorsoventral axis morphologically and functionally. Furthermore, our present study provides novel genetic model systems to investigate the crosstalk among the hippocampal complex in gene expression, morphogenesis, cell fate specification and differentiation, connectivity, functions of learning/memory and emotion/anxiety, adaptive behaviors, and the etiology of neurological diseases. Nevertheless, many enigmas, such as whether and how the abnormalities of either the dorsal or ventral hippocampus affect the characteristics of the other, remain unsolved. In addition to the excitatory lineages and circuits, interneurons and inhibitory circuits play vital roles in maintaining the plasticity and functions of the hippocampus. We also wonder whether and how defects in interneurons and inhibitory circuits could contribute to the compromised morphogenesis, connectivity, and functions of the hippocampus and the etiology of psychiatric and neurological conditions including ID, ASD, depression, anxiety, and schizophrenia.

## Materials and Methods

### Animals

*Nr2f1-floxed* (*Nr2f1^F/F^*) mice, *Nr2f2-floxed* (*Nr2f2^F/F^*) mice, *Emx1^Cre^*mice and RX^Cre^ mice (Swindell et al., 2006) (PMID: 16850473) used in the study were of the C57B6/129 mixed background. The noon of vaginal plug day was set as the embryonic day 0.5 (E0.5). Only male mice at age of 10 weeks or older were used in the Morris water maze. For other experiments, both male and female mice were used. All animal protocols were approved by the Institutional Animal Care and Use Committee (IACUC) at the Shanghai Institute of Biochemistry and Cell Biology, Chinese Academy of Sciences (Protocols: SIBCB-NAF-14-001-S308-001). All methods were performed in accordance with the relevant guidelines and regulations. Only the littermates were used for the comparison.

### Nissl staining

We used xylene to dewax paraffin sections, followed by rinsing with 100%, 95%, and 70% ethanol. The slides were stained in 0.1% Cresyl Violet solution for 25 mins. Then the sections were washed quickly in the water and differentiated in 95% ethanol. We used 100% ethanol to dehydrate the slides, followed by rinsing with the xylene solution. Finally, the neutral resin medium was used to mount the slides.

### Immunohistochemical (IHC) staining

The paraffin sections were dewaxed and rehydrated as described above for Nissl staining. The slides were boiled in 1×antigen retrieval solution (DAKO) under microwave conditions for 15 min. After cooled to room temperature (RT), the slides were incubated with 3% H2O2 for 30 min. Then, the slides were treated with blocking buffer for 60 min at RT and then incubated with the primary antibody in the hybridization buffer (10×diluted blocking buffer) overnight (O/N) at 4 °C. The next day, the tyramide signal amplification kit (TSA) (Invitrogen) was used according to the manufacturer’s protocol. After being incubated with 1%TSA blocking buffer, the sections were treated with a biotinylated secondary antibody for 60 min at RT. After being washed with 1×PBS three times, the slides were incubated with 1×HRP-conjugated streptavidin for 1 h. Next, the tyramide working solution was prepared, including the 0.15‰ H2O2 in distilled water, the 100×diluted tyramide substrate solution (tyramide-488 or tyramide-594), and the amplification buffer. The sections were incubated with the working solution for 10 min. Then the slides were counterstained with DAPI and mounted with the antifade mounting medium (Southern Biotech) (S. Feng et al., 2017; Zhang et al., 2020). Finally, the sections were observed and images were captured with a digital fluorescence microscope (Zeiss).

The following primary antibodies were used in the study: mouse anti-NR2F1 (1:1000, R&D, Cat # PP-H8132-00), mouse anti-NR2F2 (1:2000, R&D, Cat # PP-H7147-00), rabbit anti-NR2F2 (1:2000, a gift from Dr. Zhenzhong Xu, Zhejiang University, China), rabbit anti-HuB (1:500, Abcam, Cat # ab204991), rat anti-Ctip2 (1:500, Abcam, Cat # ab18465), rabbit anti-Wfs1 (1:500, ProteinTech, Cat # 11558-1-AP), goat anti-Prox1 (1:500, R&D, Cat # AF2727), rabbit anti-Calretinin (1:500, Sigma, Cat # C7479), rabbit anti-Calbindin (1:500, Swant, Cat # CB38), mouse anti-SMI312 (1:200, Covance, Cat # SMI-312R), rabbit anti-Sox2 (1:500, Affinity BioReagents, Cat # PA1-16968), rat anti-Tbr2 (1:500, Thermo Fisher, Cat # 12-4875-82), goat anti-NeuroD1 (1:200, Santa Cruz, Cat # sc-1084), goat anti-Lhx2 (1:200, Santa Cruz, Cat # sc-19344), goat anti-Lhx5 (1:200, R&D, Cat # AF6290), goat anti-β-galactosidase (LacZ) (1:400, Biogenesis, Cat # 4600-1409), mouse anti-GFAP (1:500, Sigma, Cat # G3893), rabbit anti-Nestin (1:200, Santa Cruz, Cat # sc-20978), goat anti-Dcx (1:500, Santa Cruz, Cat # sc-8066). The following secondary antibodies were used in the study: donkey anti-mouse IgG biotin-conjugated (1:400, JacksonImmuno, Cat # 715-065-150), donkey anti-rabbit IgG biotin-conjugated (1:400, JacksonImmuno, Cat # 711-065-152), donkey anti-goat IgG biotin-conjugated (1:400, JacksonImmuno, Cat # 705-066-147), donkey anti-rat IgG biotin-conjugated (1:400, JacksonImmuno, Cat # 712-065-150).

### Western blotting

We homogenized the isolated dorsal and ventral hippocampus tissues from 1-month-old mice respectively in the RIPA buffer (Applygen) with protease inhibitor cocktail (Sigma) and phosphatase inhibitors (Invitrogen) and then centrifuged at the speed of 12,000 rpm for 30 mins. We collected the supernatants and analyzed the total concentrations by the BCA kit (Applygen). Gradient SDS-PAGE gels were used to separate the same amounts of protein sample (40 µg/lane), and then the proteins were transferred to the PVDF membranes (Millipore). After being blocked by the 3% BSA (Sigma) for 2h, the membranes that contained proteins were incubated by primary antibodies at 4 °C O/N. The membranes were rinsed with 1×PBST three times for 10 mins and then treated with biotinylated secondary antibodies for 2h at RT. After washing with 1×PBST, membranes were treated with the HRP-conjugated streptavidin for 1h at RT. Finally, we used the chemiluminescence detection system (Tanon) to detect the bands. The density of the protein band was analyzed by the software Image J.

The primary antibodies were used in the experiment as below: mouse anti-NR2F1 (1:2000, R&D, Cat # PP-H8132-00), rabbit anti-NR2F2 (1:3000, a gift from Dr. Zhenzhong Xu, Zhejiang University, China), mouse anti-GAPDH (1:1000, Santa Cruz, Cat # sc-32233). The following secondary antibodies were applied in the study, including goat anti-mouse IgG biotin-conjugated (1:1000, KPL, Cat # 16-18-06), goat anti-rabbit IgG biotin-conjugated (1:1000, KPL, Cat # 16-15-06).

### Golgi staining

Deep anesthesia was performed before sacrificing the control and mutant mice, and the brains were immediately isolated. The FD rapid GolgiStain kit (FD NeuroTech) was used to process the brain tissue samples, which were immersed in an equal volume of immersion solution mixed with solutions A and B and stored in the dark for two weeks at RT. At least 5mL of immersion solution was used for each cubic meter of the tissue. To achieve the best results, the container of tissue was gently swirled from side to side twice a week during the incubating period. Afterwards, the brain tissue was transferred to solution C in the dark at RT for at least 72 hours (up to 1 week). Finally, the tissues were cut into 100 µm thick slices with a cryostat at −20°C to −22°C and transferred to gelatin-coated microscope slides containing solution C using a sample retriever. The slices were dried naturally at RT. The concrete staining procedure of the kit was followed using the manufactory’s protocol. Then, the sections were rinsed twice with double-distilled water for 4 minutes each time. The slices were placed in a mixture of one volume of solution D, one volume of solution E, and two volumes of double distilled water for 10 minutes and were rinsed twice with distilled water for 4 minutes each time. The sections were later dehydrated in 50%, 75%, and 95% ethanol for 4 minutes respectively. Next, the slices were dehydrated in 100% ethanol 4 times for 4 minutes each time. Finally, the sections were cleared in xylene and mounted with a neutral resin medium.

### Morris water maze

By recording the time spent by the mice swimming in the water tank and finding the escape platform hidden underwater, and the swimming trajectory, the Morris water maze test can objectively reflect the spatial learning and memory ability of the mice. The test was divided into the control and mutant group with at least eight mice in each group. We poured tap water into the water maze tank and added an appropriate amount of well-mixed, milky white food dye. The height of the liquid level was about 1 cm higher than the escape platform, and the water temperature was kept at about 25±1℃. At the same time, four markers of different shapes were pasted on the four directions of the inner wall above the water tank to distinguish different directions. The Morris water maze test was divided into the training phase and the probe trial phase. The training phase lasted for 6 days, 4 times a day, and the interval between each training was about 30 minutes. During training, the mice were placed into the tank from the entry points of four different quadrants facing the inner wall, and their latency was recorded from the time they entered the water to the time they found a hidden underwater platform and stood on it. After the mouse found the platform, we let it stay on the platform for 10 seconds before removing it. If the mouse failed to discover the platform 60 seconds after entering the water, it was guided to find the platform and left to stay for 10 seconds. Each mouse was placed into the water tank from four water entry points and recorded as one training session. The probe trial was carried out on the seventh day, and the underwater platform was removed. Each experimental mouse was put into the water tank at the same water entry point and allowed to move for 60 seconds. The time that each mouse spent in the quadrant, where the platform was originally placed, was recorded.

### RNA isolation and quantitative real-time PCR

Total RNAs were prepared from the whole telencephalon of the control (n=5) and double mutant (n=3) mice at E11.5 respectively, with the TRIzol Reagent (Invitrogen) by following the manufactory’s protocol. Reverse-transcription PCR and real-time quantitative PCR assays were performed as described previously (Tang et al., 2012). A student’s t-test was used to compare the means of the relative mRNA levels between the control group and mutant group. Primer sequences are as follows:

*Axin2-f*, 5’-*CTGCTGGTCAGGCAGGAG-*3’, *Axin2-r*, 5’-*TGCCAGTTTCTTTGGCTCTT-*3’; *Nr2f1-f*, 5’-*CAAAGCCATCGTGCTATTCA*-3’, *Nr2f1-r*, 5’-*CCTGCAGGCTTTCGATGT*-3’; *Nr2f2-f*, 5’-*CCTCAAAGTGGGCATGAGAC*-3’, *Nr2f2-r*, 5’-*TGGGTAGGCTGGGTAGGAG*-3’; *Emx1-f*, 5’-*CTCTCCGAGACGCAGGTG*-3’, *Emx1-r,* 5’*-CTCAGACTCCGGCCCTTC-*3’; *Emx2-f*, 5’-*CACGCTTTTGAGAAGAACCA*-3’, *Emx2-r,* 5’*-GTTCTCCGGTTCTGAAACCA-*3’; *Foxg1-f*, 5’-*GAAGGCCTCCACAGAACG*-3’, *Foxg1-r*, 5’-*GGCAAGGCATGTAGCAAAAG*-3’; *Gli3-f*, 5’-*TGATCCATCTCCTATTCCTCCA-*3’, *Gli3-r*, 5’-*TCTGGATACGTCGGGCTACT-*3’; *Lef1-f*, 5’-*TCCTGAAATCCCCACCTTCT-*3’, *Lef1-r*, 5’-*TGGGATAAACAGGCTGACCT-*3’; *Lhx2-f*, *5*’*-CAGCTTGCGCAAAAGACC-*3’, *Lhx2-r*, 5’*-TAAAAGGTTGCGCCTGAACT-*3’; *Lhx5-f*, 5’*-TGTGCAATAAGCAGCTATCCA-*3’, *Lhx5-r,* 5’*-CAAACTGCGGTCCGTACA-*3’; *Otx1-f*, 5’-*CCAGAGTCCAGAGTCCAGGT-*3’, *Otx1-r*, 5’-*CCGGGTTTTCGTTCCATT-*3’; *Otx2-f*, 5’-*GGTATGGACTTGCTGCATCC*-3’, *Otx2-r*, 5’-*CGAGCTGTGCCCTAGTAAATG*-3’; *Pax6-f*, 5’-*GTTCCCTGTCCTGTGGACTC*-3’, *Pax6-r*, 5’-*ACCGCCCTTGGTTAAAGTCT*-3’; *Tcf4-f*, 5’-*AAATGGCCACTGCTTGATGT-*3’, *Tcf4-r*, 5’-*GCACCACCGGTACTTTGTTC-*3’.

### Quantification and statistical analysis

The number of specified immunofluorescent marker-positive cells was assessed by Image J Cell Counter in full image fields. Three brain sections per mouse were counted for each index. GraphPad Prism 7.0 (GraphPad) was used to perform statistical analysis. The data analysis used one-way analysis of variance (ANOVA), Dunnett’s or Tukey’s post hoc tests, and student’s unpaired t-test. The data were expressed as the mean ± SEM. The data obtained from at least three independent replicates were used for statistical analysis. P< 0.05 was considered the significant statistical difference.

## Acknowledgements

We thank Ms. Emerald Tang for her assistance with the manuscript. This work was supported by National Natural Science Foundation of China (31671508) and Guangdong Provincial Basic and Applied Basic Research Fund (2021A1515011299) to K.T; and in part by the National Key Basic Research and Development Program of China (2018YFA0108500, 2019YFA0801402, 2018YFA0800100, 2018YFA0108000, 2018YFA0107200, 2017YFA0102700), “Strategic Priority Research Program” of the Chinese Academy of Sciences, Grant No. (XDA16020501, XDA16020404) to N. J.

## Data availability

Numerical data are available in the manuscript and supporting files.

## The information of the source data files

**Figure 1-source data 1.** The Nissl staining results of the control and RX^Cre/+^; *Nr2f2^F/F^* mutant mice at 1M (part 1); the expression of HuB and Ctip2 in the hippocampus of the control and RX^Cre/+^; *Nr2f2^F/F^* mutant mice at 1M (part 1).

**Figure 1-source data 2.** The expression of NR2F1 and NR2F2 in coronal sections and sagittal sections of the mouse brain at 1M (part 1); the expression of HuB and Ctip2 in the hippocampus of the control and RX^Cre/+^; *Nr2f2^F/F^* mutant mice at 1M (part 2).

**Figure 1-source data 3.** The expression of NR2F1 and NR2F2 in coronal sections and sagittal sections of the mouse brain at 1M (part 2).

**Figure 1-source data 4.** The expression of NR2F1 and NR2F2 in sagittal sections of the mouse brain at 1M (part 1).

**Figure 1-source data 5.** The expression of NR2F1 and NR2F2 in sagittal sections of the mouse brain at 1M (part 2); the Nissl staining results of the control and RX^Cre/+^; *Nr2f2^F/F^* mutant mice at 1M (part 2).

**Figure 1-source data 6.** The expression of NR2F1 and NR2F2 in sagittal sections of the mouse brain at 1M (part 3); Western blots data for the expression of NR2F1 and NR2F2 in the dorsal and ventral hippocampus at 1M; the Nissl staining results of the control and RX^Cre/+^; *Nr2f2^F/F^* mutant mice at 1M (part 3).

**Figure 1-figure supplement 1-source data 1.** The expression of *Nr2f1* and *Nr2f2* genes in the developing hippocampus at E12.5; the deletion efficiency of RXCre recombinase in the hippocampus of RX^Cre/+^; *Nr2f2^F/F^*, RX^Cre/+^; *Nr2f1^F/F^*, and RX^Cre/+^; *Nr2f1^F/F^*; *Nr2f2^F/F^* mice (part 1).

**Figure 1-figure supplement 1-source data 2.** The expression of *Nr2f1* and *Nr2f2* genes in the telencephalon at E10.5, E11.5, E14.5, and P0; the deletion efficiency of RXCre recombinase in the hippocampus of RX^Cre/+^; *Nr2f2^F/F^*, RX^Cre/+^; *Nr2f1^F/F^*, and RX^Cre/+^; *Nr2f1^F/F^*; *Nr2f2^F/F^* mice (part 2).

**Figure 2-source data 1.** The Nissl staining results of the control and RX^Cre/+^; *Nr2f1^F/F^* mutant mice at 1M (part 1).

**Figure 2-source data 2.** The Nissl staining results of the control and RX^Cre/+^; *Nr2f1^F/F^* mutant mice at 1M (part 2); the expression of HuB and Wfs1 in the hippocampus of the control and RX^Cre/+^; *Nr2f1^F/F^* mutant mice at 1M.

**Figure 2-source data 3.** The expression of HuB and Ctip2 in the hippocampus of the control and RX^Cre/+^; *Nr2f1^F/F^* mutant mice at 1M.

**Figure 2-source data 4.** The expression of HuB, Wfs1, and Ctip2 in the hippocampus of the control and RX^Cre/+^; *Nr2f1^F/F^* mutant mice at 1M.

**Figure 2-figure supplement 1-source data 1.** The expression of Wfs1 and Ctip2 in the dorsal hippocampus of the control and *Emx1^Cre/+^*; *Nr2f1^F/F^* mutant mice at 3M; Golgi staining results of the dorsal hippocampal CA1 pyramidal neurons and Morris water maze behavior test data of the control and *Emx1^Cre/+^*; *Nr2f1^F/F^* mutant mice.

**Figure 3-source data 1.** The Nissl staining results of the control and RX^Cre/+^; *Nr2f1^F/F^*; *Nr2f2^F/F^* double-gene mutant mice at 3W.

**Figure 3-source data 2.** The expression of HuB and Ctip2 in the hippocampus of the control and RX^Cre/+^; *Nr2f1^F/F^*; *Nr2f2^F/F^* double-gene mutant mice at 3W.

**Figure 3-source data 3.** The expression of HuB and Prox1 in the hippocampus of the control and RX^Cre/+^; *Nr2f1^F/F^*; *Nr2f2^F/F^*double-gene mutant mice at 3W.

**Figure 3-source data 4.** The expression of HuB, Ctip2, and Prox1 in the hippocampus of the control and RX^Cre/+^; *Nr2f1^F/F^*; *Nr2f2^F/F^*double-gene mutant mice at 3W.

**Figure 3-figure supplement 1-source data 1.** The expression of GFAP, Nestin, and Dcx in the SGZ of vDG in the control and RX^Cre/+^; *Nr2f2^F/F^* mutant mice at 1M, in the SGZ of dDG in the control and RX^Cre/+^; *Nr2f1^F/F^*mutant mice at 1M, and in the SGZ of DG in the control and RX^Cre/+^; *Nr2f1^F/F^*; *Nr2f2^F/F^* double-gene mutant mice at 3W (part 1).

**Figure 3-figure supplement 1-source data 2.** The expression of GFAP, Nestin, and Dcx in the SGZ of vDG in the control and RX^Cre/+^; *Nr2f2^F/F^* mutant mice at 1M, in the SGZ of dDG in the control and RX^Cre/+^; *Nr2f1^F/F^*mutant mice at 1M, and in the SGZ of DG in the control and RX^Cre/+^; *Nr2f1^F/F^*; *Nr2f2^F/F^* double-gene mutant mice at 3W (part 2).

**Figure 3-figure supplement 1-source data 3.** The expression of GFAP and Nestin in the SGZ of vDG in the control and RX^Cre/+^; *Nr2f2^F/F^*mutant mice at 1M, in the SGZ of dDG in the control and RX^Cre/+^; *Nr2f1^F/F^* mutant mice at 1M, and in the SGZ of DG in the control and RX^Cre/+^; *Nr2f1^F/F^*; *Nr2f2^F/F^*double-gene mutant mice at 3W; quantitative analysis of GFAP/Nestin-positive cells in the SGZ of DG in the control and RX^Cre/+^; *Nr2f1^F/F^*; *Nr2f2^F/F^* double-gene mutant mice at 3W.

**Figure 3-figure supplement 1-source data 4.** The expression of GFAP, Nestin, and Dcx in the SGZ of vDG in the control and RX^Cre/+^; *Nr2f2^F/F^* mutant mice at 1M, in the SGZ of dDG in the control and RX^Cre/+^; *Nr2f1^F/F^*mutant mice at 1M, and in the SGZ of DG in the control and RX^Cre/+^; *Nr2f1^F/F^*; *Nr2f2^F/F^* double-gene mutant mice at 3W (part 3); quantitative analysis of Dcx-positive cells in the SGZ of DG in the control and RX^Cre/+^; *Nr2f1^F/F^*; *Nr2f2^F/F^* double-gene mutant mice at 3W.

**Figure 4-source data 1.** The expression of Calretinin, Calbindin, and SMI312 in the ventral hippocampus of the control and RX^Cre/+^; *Nr2f2^F/F^* single-gene mutant mice at 1M.

**Figure 4-source data 2.** The expression SMI312 in the dorsal hippocampus of the control and RX^Cre/+^; *Nr2f1^F/F^* single-gene mutant mice at 1M, and the expression of Calretinin and Calbindin in the hippocampus of the control and RX^Cre/+^; *Nr2f1^F/F^*; *Nr2f2^F/F^*double-gene mutant mice at 3W.

**Figure 4-source data 3.** The expression of Calretinin and Calbindin in the dorsal hippocampus of the control and RX^Cre/+^; *Nr2f1^F/F^* single-gene mutant mice at 1M, and the expression of

SMI312 in the hippocampus of the control and RX^Cre/+^; *Nr2f1^F/F^*; *Nr2f2^F/F^* double-gene mutant mice at 3W.

**Figure 5-source data 1.** The expression of Lhx5 and Lhx2 in the telencephalon of the control and RX^Cre/+^; *Nr2f1^F/F^*; *Nr2f2^F/F^* double-mutant mice at E14.5; the expression of Tbr2 and NeuroD1 in the telencephalon of the control and RX^Cre/+^; *Nr2f1^F/F^*; *Nr2f2^F/F^* double-mutant mice at E14.5 (part 1).

**Figure 5-source data 2.** The expression of Sox2, Tbr2, and NeuroD1 in the telencephalon of the control and RX^Cre/+^; *Nr2f1^F/F^*; *Nr2f2^F/F^*double-mutant mice at E14.5.

**Figure 5-source data 3.** The expression profiles of genes involved in the hippocampal development of the control and double mutant mice at E11.5; the expression of Lhx5 and Lhx2 in the telencephalon of the control and RX^Cre/+^; *Nr2f1^F/F^*; *Nr2f2^F/F^* double-mutant mice at E11.5 and E13.5; the expression of Tbr2 and NeuroD1 in the telencephalon of the control and RX^Cre/+^; *Nr2f1^F/F^*; *Nr2f2^F/F^* double-mutant mice at E14.5 (part 2); quantitative analysis of Tbr2-positive and NeuroD1-positive cells in the hippocampal primordium of the control and RX^Cre/+^; *Nr2f1^F/F^*; *Nr2f2^F/F^* double-mutant mice at E14.5.

**Figure 5-figure supplement 1-source data 1.** Roles of *Nr2f1* and *Nr2f2* genes in the development and function of the hippocampus and the association with neurological disorders.

---

**Figure 1—figure supplement 1.**
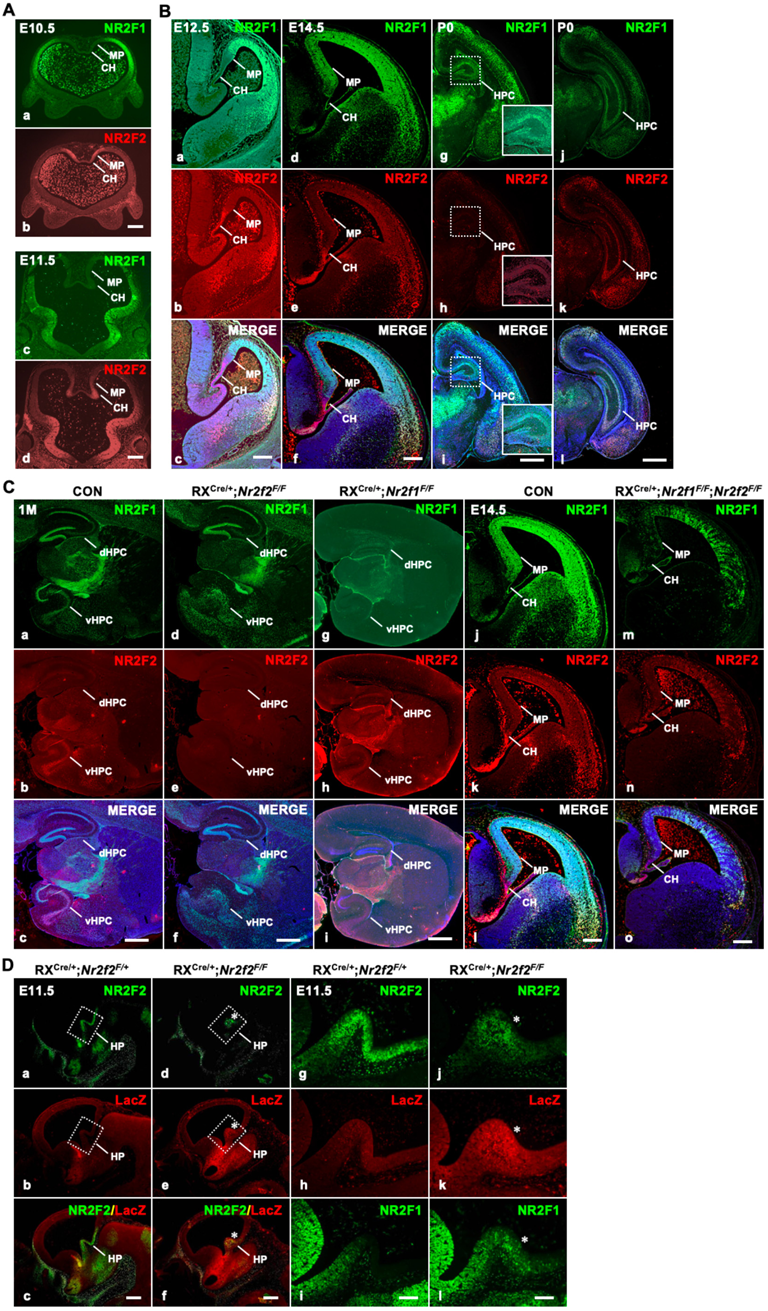
The expression of *Nr2f* genes in the early developing hippocampus and different conditional knock mouse models. A, The expression of *Nr2f1* and *Nr2f2* genes in the forebrain at E10.5 (**a-b**) and E11.5 (**c-d**). **B,** The expression of *Nr2f1* and *Nr2f2* genes in the developing hippocampus at E12.5 (**a-c**), E14.5 (**d-f**), and P0 (**g-l**). **C,** Compared with that of control mice (**a-c**), *Nr2f2* is efficiently deleted by RXCre recombinase in the hippocampus of *Nr2f2* mutant mice at 1M (**d-f**); *Nr2f1* is clearly deleted by RXCre recombinase in the hippocampus of *Nr2f1* mutant mice at 1M (**g-i**). Compared with that of control mice (**j-l**), NR2F1 and NR2F2 were efficiently deleted by RXCre recombinase at the hippocampal primordium, including the MP and CH, in *Nr2f1/2* double-mutant mice at E14.5 (**m-o**). **D,** Compared with that of the control mice (**a-c, g-h**), the expression of *Nr2f2* was significantly decreased in the hippocampal primordium of the homozygous mutant mice at E11.5; meanwhile, the LacZ signals obviously increased in the *Nr2f2* homozygous mutant mice at E11.5 (**d**-**f, j-k**). Compared with that of the control mice (**i**), the expression of NR2F1 is activated in the caudal hippocampal primordium of the homozygous mutant mice at E11.5 (**l**). CH, cortical hem; dHPC, dorsal hippocampus; HP, hippocampal primordium; HPC, hippocampus; MP, medial pallium; vHPC, ventral hippocampus. Scale bars, **Aa-d**, **Ba-f**, **Cj-o**, 200 μm; **Bg-l**, **Ca-i**, **Dg-l**, 100 μm; **Da-f**, 250 μm.

**Figure 2—figure supplement 1.**
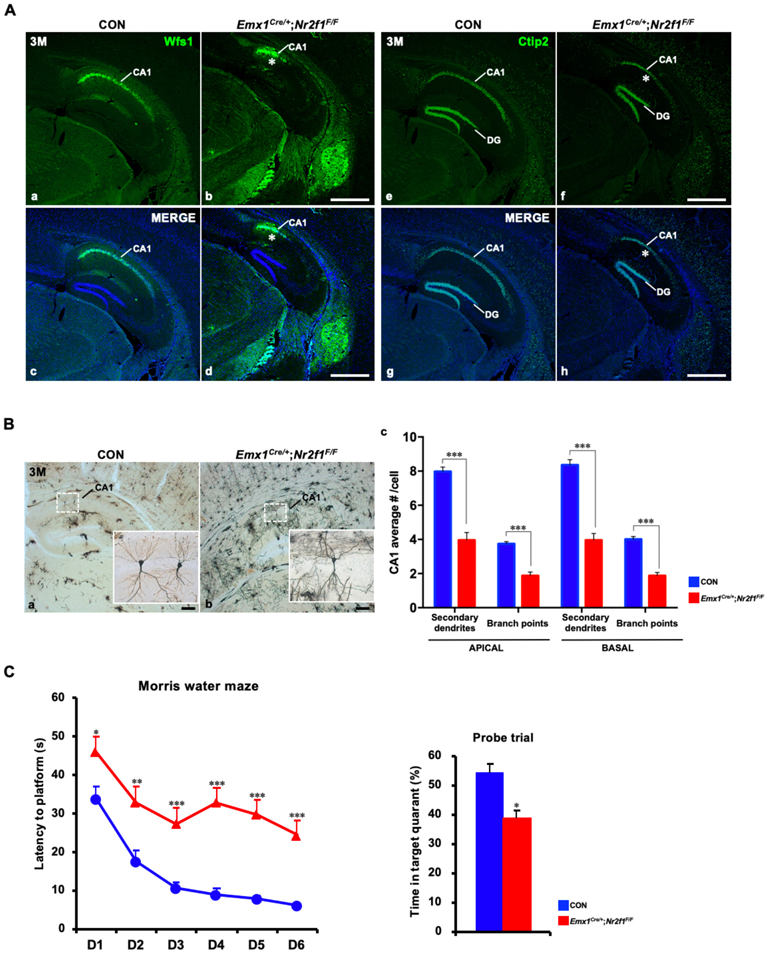
Defects in *Emx1^Cre/+^*; *Nr2f1^F/F^* mutant mice. **A**, Immunofluorescence staining data showed that compared with those of control mice (**a, c, e, g**), the proportions of either the Wfs1- or Ctip2-positive dorsal CA1 domain were reduced in *Emx1^Cre/+^*; *Nr2f1^F/F^* mutant mice at 3M (**b, d, f, h**). **B,** Golgi staining showed that compared with those of controls, the numbers of branch points and secondary dendrites of both apical and basal dendrites were significantly reduced in the dorsal hippocampal CA1 pyramidal neurons of *Emx1^Cre/+^*; *Nr2f1^F/F^* mutant mice (**a-c**). **C,** The Morris water maze behavior test showed that compared with that of controls, spatial learning and memory were significantly damaged in *Emx1^Cre/+^*; *Nr2f1^F/F^* mutant mice. Data are expressed as the mean ± SEM. *Student’s t test*, \**P*<0.05, \*\**P*<0.01, \*\*\**P*<0.001. Scale bars, **Aa-h,** 100 μm; **Ba-b,** 50 μm.

**Figure 3—figure supplement 1.**
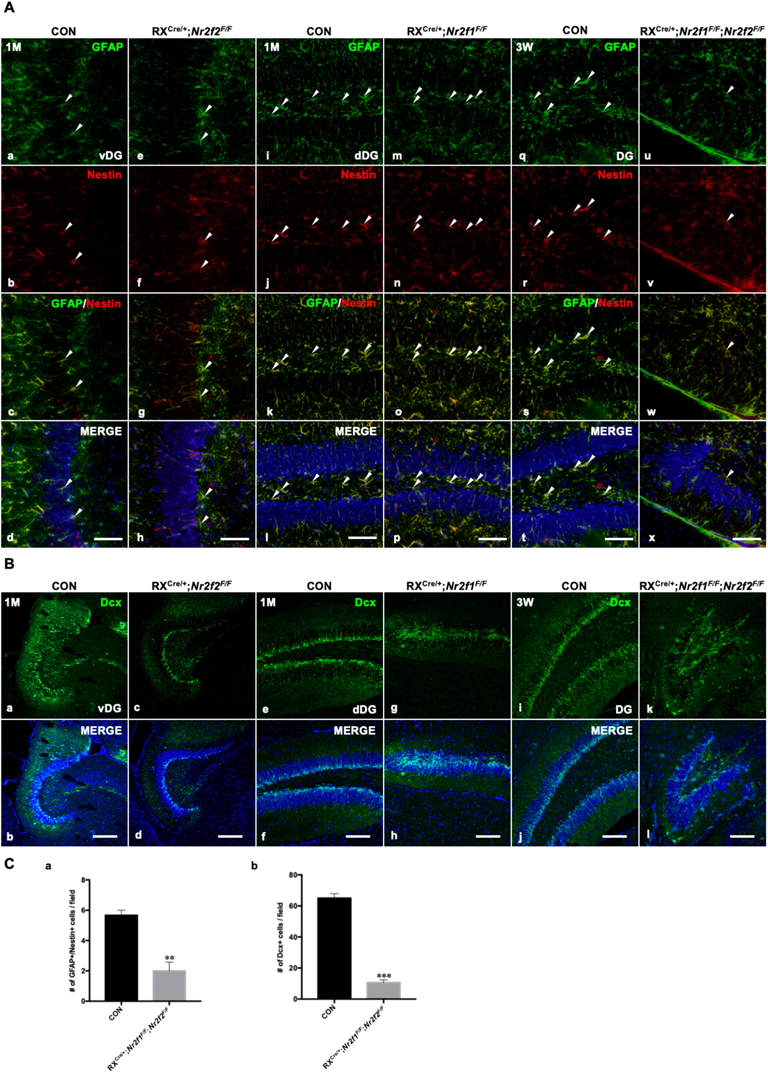
Adult neurogenesis was abnormal in the hippocampi of *Nr2f1/2* double-gene mutant mice. **A,** The expression of GFAP and Nestin, markers of NSCs, in the SGZ of the vDG in control and *Nr2f2* mutant mice at 1M (**a-h**), in the SGZ of the dDG in control and *Nr2f1* mutant mice at 1M (**i-p**) and in the SGZ of the DG in control and *Nr2f1/2* double-gene mutant mice at 3W (**q-x**). **B**, The expression of Dcx, a marker of newborn neurons, in the SGZ of the vDG in control and *Nr2f2* mutant mice at 1M (**a-d**), in the SGZ of the dDG in control and *Nr2f1* mutant mice at 1M (**e-h**) and in the SGZ of the DG in control and *Nr2f1/2* double-gene mutant mice at 3W (**i-l**). **C,** Quantitative analysis of GFAP/Nestin-positive cells (**a**) and Dcx-positive cells (**b**) in the SGZ of the DG in control and *Nr2f1/2* double-gene mutant mice at 3W. The numbers of GFAP and Nestin double-positive NSCs and Dcx-positive newborn neurons were significantly reduced in double-mutant mice. Data are expressed as the mean ± SEM. *Student’s t test*, \*\**P*<0.01, \*\*\**P*<0.001. dDG, dorsal DG; NSC, neural stem cell; SGZ, subgranular zone; vDG, ventral DG. Scale bars, **Aa-x**, 50 μm; **Ba-l**, 100 μm.

**Figure 5—figure supplement 1.**
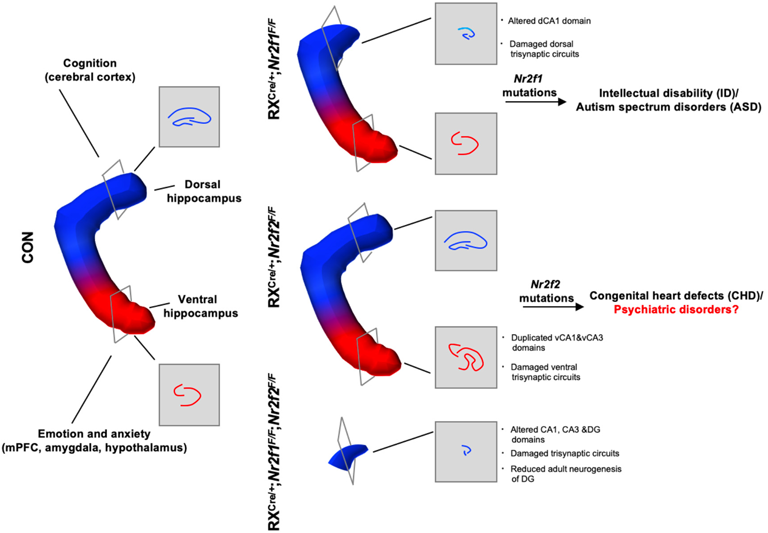
*Nr2f1* and *Nr2f2* genes coordinate to control distinct characteristics of the hippocampus. Roles of *Nr2f1* and *Nr2f2* genes in the development and function of the hippocampus and the association with neurological diseases. *Nr2f1* is required for the morphogenesis of the dorsal hippocampus and the specification of dorsal CA1 pyramidal neuron lineage, which are associated with neurodevelopmental disorders, including ID and ASD. *Nr2f2* is required to prevent the duplication of the CA1 and CA3 lineages of the ventral hippocampus, which may be related to psychiatric diseases such as depression, anxiety, or schizophrenia. The *Nr2f1* and *Nr2f2* genes are novel intrinsic regulatory genes, which cooperate with each other to ensure the early morphogenesis of the hippocampus.

## References

Al Turki, S., Manickaraj, A. K., Mercer, C. L., Gerety, S. S., Hitz, M. P., Lindsay, S., … Hurles, M. E. (2014). Rare variants in NR2F2 cause congenital heart defects in humans. Am J Hum Genet, 94(4), 574–585. doi:10.1016/j.ajhg.2014.03.007

Alzu’bi, A., Lindsay, S. J., Harkin, L. F., McIntyre, J., Lisgo, S. N., & Clowry, G. J. (2017). The Transcription Factors COUP-TFI and COUP-TFII have Distinct Roles in Arealisation and GABAergic Interneuron Specification in the Early Human Fetal Telencephalon. Cereb Cortex, 27(10), 4971–4987. doi:10.1093/cercor/bhx185

Amaral, D. G. (1993). Emerging principles of intrinsic hippocampal organization. Current Opinion in Neurobiology, 3(2), 225–229. 10.1016/0959-4388(93)90214-J

Anacker, C., & Hen, R. (2017). Adult hippocampal neurogenesis and cognitive flexibility - linking memory and mood. Nat Rev Neurosci, 18(6), 335–346. doi:10.1038/nrn.2017.45

Armentano, M., Chou, S. J., Tomassy, G. S., Leingärtner, A., O’Leary, D. D., & Studer, M. (2007). COUP-TFI regulates the balance of cortical patterning between frontal/motor and sensory areas. Nat Neurosci, 10(10), 1277–1286. doi:10.1038/nn1958

Baik, J. H. (2020). Stress and the dopaminergic reward system. Exp Mol Med, 52(12), 1879–1890. doi:10.1038/s12276-020-00532-4

Bast, T. (2007). Toward an integrative perspective on hippocampal function: from the rapid encoding of experience to adaptive behavior. Rev Neurosci, 18(3-4), 253–281. doi:10.1515/revneuro.2007.18.3-4.253

Basu, J., & Siegelbaum, S. A. (2015). The Corticohippocampal Circuit, Synaptic Plasticity, and Memory. Cold Spring Harb Perspect Biol, 7(11). doi:10.1101/cshperspect.a021733

Bertacchi, M., Romano, A. L., Loubat, A., Tran Mau-Them, F., Willems, M., Faivre, L., … Studer, M. (2020). NR2F1 regulates regional progenitor dynamics in the mouse neocortex and cortical gyrification in BBSOAS patients. EMBO J, 39(13), e104163. doi:10.15252/embj.2019104163

Bliss, T. V., & Lomo, T. (1973). Long-lasting potentiation of synaptic transmission in the dentate area of the anaesthetized rabbit following stimulation of the perforant path. J Physiol, 232(2), 331–356. doi:10.1113/jphysiol.1973.sp010273

Bosch, D. G., Boonstra, F. N., Gonzaga-Jauregui, C., Xu, M., de Ligt, J., Jhangiani, S., … Schaaf, C. P. (2014). NR2F1 mutations cause optic atrophy with intellectual disability. Am J Hum Genet, 94(2), 303–309. doi:10.1016/j.ajhg.2014.01.002

Bryant, K. G., & Barker, J. M. (2020). Arbitration of Approach-Avoidance Conflict by Ventral Hippocampus. Front Neurosci, 14, 615337. doi:10.3389/fnins.2020.615337

Cenquizca, L. A., & Swanson, L. W. (2007). Spatial organization of direct hippocampal field CA1 axonal projections to the rest of the cerebral cortex. Brain Res Rev, 56(1), 1–26. doi:10.1016/j.brainresrev.2007.05.002

Contesse, T., Ayrault, M., Mantegazza, M., Studer, M., & Deschaux, O. (2019). Hyperactive and anxiolytic-like behaviors result from loss of COUP-TFI/Nr2f1 in the mouse cortex. Genes Brain Behav, 18(7), e12556. doi:10.1111/gbb.12556

Del Pino, I., Tocco, C., Magrinelli, E., Marcantoni, A., Ferraguto, C., Tomagra, G., … Studer, M. (2020). COUP-TFI/Nr2f1 Orchestrates Intrinsic Neuronal Activity during Development of the Somatosensory Cortex. Cereb Cortex, 30(11), 5667–5685. doi:10.1093/cercor/bhaa137

Eichenbaum, H. (2017). On the Integration of Space, Time, and Memory. Neuron, 95(5), 1007–1018. doi:10.1016/j.neuron.2017.06.036

Eichenbaum, H., & Cohen, N. J. (2014). Can we reconcile the declarative memory and spatial navigation views on hippocampal function? Neuron, 83(4), 764–770. doi:10.1016/j.neuron.2014.07.032

Fanselow, M. S., & Dong, H. W. (2010). Are the dorsal and ventral hippocampus functionally distinct structures? Neuron, 65(1), 7–19. doi:10.1016/j.neuron.2009.11.031

Feng, J., Hsu, W. H., Patterson, D., Tseng, C. S., Hsing, H. W., Zhuang, Z. H., … Chou, S. J. (2021). COUP-TFI specifies the medial entorhinal cortex identity and induces differential cell adhesion to determine the integrity of its boundary with neocortex. Sci Adv, 7(27). doi:10.1126/sciadv.abf6808

Feng, S., Xing, C., Shen, T., Qiao, Y., Wang, R., Chen, J., … Tang, K. (2017). Abnormal Paraventricular Nucleus of Hypothalamus and Growth Retardation Associated with Loss of Nuclear Receptor Gene COUP-TFII. Sci Rep, 7(1), 5282. doi:10.1038/s41598-017-05682-6

Flore, G., Di Ruberto, G., Parisot, J., Sannino, S., Russo, F., Illingworth, E. A., … De Leonibus, E. (2017). Gradient COUP-TFI Expression Is Required for Functional Organization of the Hippocampal Septo-Temporal Longitudinal Axis. Cereb Cortex, 27(2), 1629–1643. doi:10.1093/cercor/bhv336

Galceran, J., Miyashita-Lin, E. M., Devaney, E., Rubenstein, J. L., & Grosschedl, R. (2000). Hippocampus development and generation of dentate gyrus granule cells is regulated by LEF1. Development, 127(3), 469–482. doi:10.1242/dev.127.3.469

Gao, X., Arlotta, P., Macklis, J. D., & Chen, J. (2007). Conditional knock-out of beta-catenin in postnatal-born dentate gyrus granule neurons results in dendritic malformation. J Neurosci, 27(52), 14317–14325. doi:10.1523/jneurosci.3206-07.2007

Gould, E., & Cameron, H. A. (1996). Regulation of neuronal birth, migration and death in the rat dentate gyrus. Dev Neurosci, 18(1-2), 22–35. doi:10.1159/000111392

Hebert, J. M., & Fishell, G. (2008). The genetics of early telencephalon patterning: some assembly required. Nat Rev Neurosci, 9(9), 678–685. doi:10.1038/nrn2463

Herman, J. P., McKlveen, J. M., Ghosal, S., Kopp, B., Wulsin, A., Makinson, R., … Myers, B. (2016). Regulation of the Hypothalamic-Pituitary-Adrenocortical Stress Response. Compr Physiol, 6(2), 603–621. doi:10.1002/cphy.c150015

High, F. A., Bhayani, P., Wilson, J. M., Bult, C. J., Donahoe, P. K., & Longoni, M. (2016). De novo frameshift mutation in COUP-TFII (NR2F2) in human congenital diaphragmatic hernia. Am J Med Genet A, 170(9), 2457–2461. doi:10.1002/ajmg.a.37830

Hoover, W. B., & Vertes, R. P. (2007). Anatomical analysis of afferent projections to the medial prefrontal cortex in the rat. Brain Struct Funct, 212(2), 149–179. doi:10.1007/s00429-007-0150-4

Kandel, E. R., Dudai, Y., & Mayford, M. R. (2014). The molecular and systems biology of memory. Cell, 157(1), 163–186. doi:10.1016/j.cell.2014.03.001

Kim, B. J., Takamoto, N., Yan, J., Tsai, S. Y., & Tsai, M. J. (2009). Chicken Ovalbumin Upstream Promoter-Transcription Factor II (COUP-TFII) regulates growth and patterning of the postnatal mouse cerebellum. Dev Biol, 326(2), 378–391. doi:10.1016/j.ydbio.2008.11.001

Kishi, T., Tsumori, T., Ono, K., Yokota, S., Ishino, H., & Yasui, Y. (2000). Topographical organization of projections from the subiculum to the hypothalamus in the rat. J Comp Neurol, 419(2), 205–222. doi:10.1002/(sici)1096-9861(20000403)419:2<205::aid-cne5>3.0.co;2-0

Lee, H., GoodSmith, D., & Knierim, J. J. (2020). Parallel processing streams in the hippocampus. Curr Opin Neurobiol, 64, 127–134. doi:10.1016/j.conb.2020.03.004

Lee, S. M., Tole, S., Grove, E., & McMahon, A. P. (2000). A local Wnt-3a signal is required for development of the mammalian hippocampus. Development, 127(3), 457–467. doi:10.1242/dev.127.3.457

Leid, M., Ishmael, J. E., Avram, D., Shepherd, D., Fraulob, V., & Dolle, P. (2004). CTIP1 and CTIP2 are differentially expressed during mouse embryogenesis. Gene Expr Patterns, 4(6), 733–739. doi:10.1016/j.modgep.2004.03.009

Li, J., Sun, L., Peng, X. L., Yu, X. M., Qi, S. J., Lu, Z. J., … Shen, Q. (2021). Integrative genomic analysis of early neurogenesis reveals a temporal genetic program for differentiation and specification of preplate and Cajal-Retzius neurons. PLoS Genet, 17(3), e1009355. doi:10.1371/journal.pgen.1009355

Lisman, J., Buzsaki, G., Eichenbaum, H., Nadel, L., Ranganath, C., & Redish, A. D. (2017). Viewpoints: how the hippocampus contributes to memory, navigation and cognition. Nat Neurosci, 20(11), 1434–1447. doi:10.1038/nn.4661

Lodato, S., Tomassy, G. S., De Leonibus, E., Uzcategui, Y. G., Andolfi, G., Armentano, M., … Studer, M. (2011). Loss of COUP-TFI alters the balance between caudal ganglionic eminence- and medial ganglionic eminence-derived cortical interneurons and results in resistance to epilepsy. J Neurosci, 31(12), 4650–4662. doi:10.1523/jneurosci.6580-10.2011

Mangale, V. S., Hirokawa, K. E., Satyaki, P. R., Gokulchandran, N., Chikbire, S., Subramanian, L., … Monuki, E. S. (2008). Lhx2 selector activity specifies cortical identity and suppresses hippocampal organizer fate. Science, 319(5861), 304–309. doi:10.1126/science.1151695

Miquelajáuregui, A., Varela-Echavarría, A., Ceci, M. L., García-Moreno, F., Ricaño, I., Hoang, K., … Zhao, Y. (2010). LIM-homeobox gene Lhx5 is required for normal development of Cajal-Retzius cells. J Neurosci, 30(31), 10551–10562. doi:10.1523/jneurosci.5563-09.2010

Monuki, E. S., Porter, F. D., & Walsh, C. A. (2001). Patterning of the dorsal telencephalon and cerebral cortex by a roof plate-Lhx2 pathway. Neuron, 32(4), 591–604. doi:10.1016/s0896-6273(01)00504-9

Moore, S. A., & Iulianella, A. (2021). Development of the mammalian cortical hem and its derivatives: the choroid plexus, Cajal-Retzius cells and hippocampus. Open Biol, 11(5), 210042. doi:10.1098/rsob.210042

Moser, M. B., & Moser, E. I. (1998). Functional differentiation in the hippocampus. Hippocampus, 8(6), 608–619. doi:10.1002/(SICI)1098-1063(1998)8:6<608::AID-HIPO3>3.0.CO;2-7

O’Keefe, J., & Conway, D. H. (1978). Hippocampal place units in the freely moving rat: why they fire where they fire. Exp Brain Res, 31(4), 573–590. doi:10.1007/BF00239813

O’Keefe, J., & Dostrovsky, J. (1971). The hippocampus as a spatial map. Preliminary evidence from unit activity in the freely-moving rat. Brain Res, 34(1), 171–175. doi:10.1016/0006-8993(71)90358-1

O’Leary, O. F., & Cryan, J. F. (2014). A ventral view on antidepressant action: roles for adult hippocampal neurogenesis along the dorsoventral axis. Trends Pharmacol Sci, 35(12), 675–687. doi:10.1016/j.tips.2014.09.011

Penfield, W., & Milner, B. (1958). Memory deficit produced by bilateral lesions in the hippocampal zone. AMA Arch Neurol Psychiatry, 79(5), 475–497. doi:10.1001/archneurpsyc.1958.02340050003001

Pitkanen, A., Pikkarainen, M., Nurminen, N., & Ylinen, A. (2000). Reciprocal connections between the amygdala and the hippocampal formation, perirhinal cortex, and postrhinal cortex in rat. A review. Ann N Y Acad Sci, 911, 369–391. doi:10.1111/j.1749-6632.2000.tb06738.x

Porter, F. D., Drago, J., Xu, Y., Cheema, S. S., Wassif, C., Huang, S. P., … Westphal, H. (1997). Lhx2, a LIM homeobox gene, is required for eye, forebrain, and definitive erythrocyte development. Development, 124(15), 2935–2944. doi:10.1242/dev.124.15.2935

Roberts, A. C., Tomic, D. L., Parkinson, C. H., Roeling, T. A., Cutter, D. J., Robbins, T. W., & Everitt, B. J. (2007). Forebrain connectivity of the prefrontal cortex in the marmoset monkey (Callithrix jacchus): an anterograde and retrograde tract-tracing study. J Comp Neurol, 502(1), 86–112. doi:10.1002/cne.21300

Roy, A., Gonzalez-Gomez, M., Pierani, A., Meyer, G., & Tole, S. (2014). Lhx2 regulates the development of the forebrain hem system. Cereb Cortex, 24(5), 1361–1372. doi:10.1093/cercor/bhs421

Schuurmans, C., & Guillemot, F. (2002). Molecular mechanisms underlying cell fate specification in the developing telencephalon. Curr Opin Neurobiol, 12(1), 26–34. doi:10.1016/s0959-4388(02)00286-6

Scoville, W. B., & Milner, B. (1957). Loss of recent memory after bilateral hippocampal lesions. J Neurol Neurosurg Psychiatry, 20(1), 11–21. doi:10.1136/jnnp.20.1.11

Simeone, A., Gulisano, M., Acampora, D., Stornaiuolo, A., Rambaldi, M., & Boncinelli, E. (1992). Two vertebrate homeobox genes related to the Drosophila empty spiracles gene are expressed in the embryonic cerebral cortex. EMBO J, 11(7), 2541–2550. doi:10.1002/j.1460-2075.1992.tb05319.x

Strange, B. A., Witter, M. P., Lein, E. S., & Moser, E. I. (2014). Functional organization of the hippocampal longitudinal axis. Nat Rev Neurosci, 15(10), 655–669. doi:10.1038/nrn3785

Sugiyama, T., Osumi, N., & Katsuyama, Y. (2014). A novel cell migratory zone in the developing hippocampal formation. J Comp Neurol, 522(15), 3520–3538. doi:10.1002/cne.23621

Swindell, E. C., Bailey, T. J., Loosli, F., Liu, C., Amaya-Manzanares, F., Mahon, K. A., … Jamrich, M. (2006). Rx-Cre, a tool for inactivation of gene expression in the developing retina. Genesis, 44(8), 361–363. doi:10.1002/dvg.20225

Takeda, K., Inoue, H., Tanizawa, Y., Matsuzaki, Y., Oba, J., Watanabe, Y., … Oka, Y. (2001). WFS1 (Wolfram syndrome 1) gene product: predominant subcellular localization to endoplasmic reticulum in cultured cells and neuronal expression in rat brain. Hum Mol Genet, 10(5), 477–484. doi:10.1093/hmg/10.5.477

Tang, K., Rubenstein, J. L., Tsai, S. Y., & Tsai, M. J. (2012). COUP-TFII controls amygdala patterning by regulating neuropilin expression. Development, 139(9), 1630–1639. doi:10.1242/dev.075564

Toda, T., Parylak, S. L., Linker, S. B., & Gage, F. H. (2019). The role of adult hippocampal neurogenesis in brain health and disease. Mol Psychiatry, 24(1), 67–87. doi:10.1038/s41380-018-0036-2

Tole, S., Goudreau, G., Assimacopoulos, S., & Grove, E. A. (2000). Emx2 is required for growth of the hippocampus but not for hippocampal field specification. J Neurosci, 20(7), 2618–2625. Retrieved from https://www.ncbi.nlm.nih.gov/pubmed/10729342

Tuncdemir, S. N., Lacefield, C. O., & Hen, R. (2019). Contributions of adult neurogenesis to dentate gyrus network activity and computations. Behav Brain Res, 374, 112112. doi:10.1016/j.bbr.2019.112112

Tyng, C. M., Amin, H. U., Saad, M. N. M., & Malik, A. S. (2017). The Influences of Emotion on Learning and Memory. Front Psychol, 8, 1454. doi:10.3389/fpsyg.2017.01454

Vorhees, C. V., & Williams, M. T. (2006). Morris water maze: procedures for assessing spatial and related forms of learning and memory. Nat Protoc, 1(2), 848–858. doi:10.1038/nprot.2006.116

Yang, X., Feng, S., & Tang, K. (2017). COUP-TF Genes, Human Diseases, and the Development of the Central Nervous System in Murine Models. Curr Top Dev Biol, 125, 275–301. doi:10.1016/bs.ctdb.2016.12.002

Yoshida, M., Suda, Y., Matsuo, I., Miyamoto, N., Takeda, N., Kuratani, S., & Aizawa, S. (1997). Emx1 and Emx2 functions in development of dorsal telencephalon. Development, 124(1), 101–111. doi:10.1242/dev.124.1.101

Yu, D. X., Marchetto, M. C., & Gage, F. H. (2014). How to make a hippocampal dentate gyrus granule neuron. Development, 141(12), 2366–2375. doi:10.1242/dev.096776

Zhang, K., Yu, F., Zhu, J., Han, S., Chen, J., Wu, X., … Tang, K. (2020). Imbalance of Excitatory/Inhibitory Neuron Differentiation in Neurodevelopmental Disorders with an NR2F1 Point Mutation. Cell Rep, 31(3), 107521. doi:10.1016/j.celrep.2020.03.085

Zhao, Y., Sheng, H. Z., Amini, R., Grinberg, A., Lee, E., Huang, S., … Westphal, H. (1999). Control of hippocampal morphogenesis and neuronal differentiation by the LIM homeobox gene Lhx5. Science, 284(5417), 1155–1158. doi:10.1126/science.284.5417.1155

Zhou, C., Qiu, Y., Pereira, F. A., Crair, M. C., Tsai, S. Y., & Tsai, M. J. (1999). The nuclear orphan receptor COUP-TFI is required for differentiation of subplate neurons and guidance of thalamocortical axons. Neuron, 24(4), 847–859. doi:10.1016/s0896-6273(00)81032-6

Zhou, C., Tsai, S. Y., & Tsai, M. J. (2001). COUP-TFI: an intrinsic factor for early regionalization of the neocortex. Genes Dev, 15(16), 2054–2059. doi:10.1101/gad.913601

Zhou, X., Liu, F., Tian, M., Xu, Z., Liang, Q., Wang, C., … Yang, Z. (2015). Transcription factors COUP-TFI and COUP-TFII are required for the production of granule cells in the mouse olfactory bulb. Development, 142(9), 1593–1605. doi:10.1242/dev.115279

